# A new universal system of tree shape indices

**DOI:** 10.1101/2023.07.17.549219

**Authors:** Robert Noble, Kimberley Verity

## Abstract

The comparison and categorization of tree diagrams is fundamental to large parts of biology, linguistics, computer science, and other fields, yet the indices currently applied to describing tree shape have important flaws that complicate their interpretation and limit their scope. Here we introduce a new system of indices with no such shortcomings. Our indices account for node sizes and branch lengths and are robust to small changes in either attribute. Unlike currently popular phylogenetic diversity, phylogenetic entropy, and tree balance indices, our definitions assign interpretable values to all rooted trees and enable meaningful comparison of any pair of trees. Our self-consistent definitions further unite measures of diversity, richness, balance, symmetry, effective height, effective outdegree, and effective branch count in a coherent system, and we derive numerous simple relationships between these indices. The main practical advantages of our indices are in 1) quantifying diversity in non-ultrametric trees; 2) assessing the balance of trees that have non-uniform branch lengths or node sizes; 3) comparing the balance of trees with different leaf counts or outdegrees; 4) obtaining a coherent, generic, multidimensional quantification of tree shape that is robust to sampling error and inferential error. We illustrate these features by comparing the shapes of trees representing the evolution of HIV and of Uralic languages, and trees generated by computational models of tumour evolution. Given the ubiquity of tree structures, we identify a wide range of applications across diverse domains. tree indices, tree shape, tree balance, phylogenetic diversity, phylogenetic entropy, rooted trees

---

Tree shape indices that quantify key properties of rooted trees – such as the effective number of leaves, average out-degree, and balance – have myriad applications. Conservation biologists use phylogenetic diversity values to determine which actions will preserve the most biodiversity (Tucker et al., 2017; Veron et al., 2019). Tree balance indices are used to compare models and to infer parameter values in systematic biology (Mooers and Heard, 1997; Purvis and Agapow, 2002), virology (Chindelevitch et al., 2021; Barzilai and Schrago, 2023), epidemiology (Leventhal et al., 2012; Colijn and Gardy, 2014), and oncology (Scott et al., 2020; Noble et al., 2022). Computer scientists seek to balance binary trees to make them more efficient as data structures (Albers and Westbrook, 2005). Numerous indices designed for such tasks have previously been proposed (Pavoine and Bonsall, 2011; Tucker et al., 2017; Fischer et al., 2023). However, no existing index provides a general purpose method for fairly evaluating the shape of any rooted tree. This paper introduces a system of such indices.

Rather than simply adding to a profusion of indices, our aim here is to solve important open problems: How can we modify existing phylogenetic diversity and entropy indices so that they are meaningful when applied to non-ultrametric trees? How can we define a tree balance index that accounts for both branch lengths and node sizes? How can we likewise generalize the concepts of outdegree, branch count, and node count? How can we unite all these types of tree shape index in a coherent system, so that their interrelationships can be easily understood? Only by solving these problems can we arrive at a general purpose method for fairly evaluating the shape of any rooted tree.

Among current diversity indices for generic rooted trees, arguably the most sophisticated are those introduced by Chao et al. (2010), which generalize and unify previous definitions of Hill (1973), Faith (1992), Jost (2006) and Allen et al. (2009). In quantifying the effective number of types in a data set, tthe 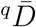 indices of Chao et al. (2010) account for both node sizes (type frequencies) and branch lengths (degree of dissimilarity between types). Nevertheless, a critical shortcoming of these indices, which limits their applications, is that they assign meaningful values only to leafy ultrametric trees (that is, trees in which the only non-zero-sized nodes are leaves, all equally distant from the root) (Chao et al., 2010; Leinster and Cobbold, 2012). We will further show that the 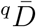 indices of Chao et al. (2010) are not fully self-consistent and have peculiar properties for *q >* 1. Moreover, the relationships between these diversity indices and other types of index, such as tree balance indices, are generally opaque, which thwarts multi-dimensional analysis.

Conventional tree balance and imbalance indices – including those attributed to Sackin (1972) and Colless (1982), the total cophenetic index of Mir et al. (2013), and others reviewed by Fischer et al. (2023) – are also flawed. These indices, which are meant to quantify the extent to which each internal node splits its descendants into equally sized subtrees, are not defined for all rooted trees, do not permit meaningful comparison of trees with differing leaf counts, and are highly sensitive to the addition or removal of rare types (Noble et al., 2022; Lemant et al., 2022). We recently introduced a family of tree balance indices that solve these problems and that have additional desirable properties (Lemant et al., 2022). Our previous definitions are defined for any degree distribution, account for node sizes, and enable meaningful comparison of trees with different numbers of leaves. But our previous definitions do not account for branch lengths, which restricts their applications because branch lengths often convey important information (for example, genetic distance in virus evolution, or elapsed time in the evolution of species).

Here we define a new system of indices that resolve all the aforementioned problems by accounting for node sizes and branch lengths, being robust to small changes to the tree, assigning meaningful values to all rooted trees, and belonging to a coherent framework, so that mathematical relationships between the indices are well characterized. Our system captures fundamental properties such as diversity (effective number of leaves), tree balance (the extent to which each internal node splits its descendants into equally sized subtrees), and bushiness (average effective outdegree). Given that our indices share the desirable properties but not the flaws of prior indices, we discuss their potential to supersede current methods in a wide range of applications.

## Materials AND Methods

### Hill numbers as a basis for defining robust, universal, interpretable tree indices

A rooted tree is a tree in which one node is designated the root and all branches are directed away from the root. Our aim is to define indices that are useful for categorizing and comparing the shapes of unlabelled rooted trees that have three attributes: tree topology, non-negative node sizes, and non-negative branch lengths. These indices should be generic and model-agnostic, meaning that they make no assumptions about what the tree represents or the process by which it was generated. In evolutionary trees, for example, the size of a node can correspond to the population size of the respective biological type, or simply to whether a type is extant (node size 1) or extinct (0), while branch lengths can represent genetic distance, morphological difference, or elapsed time.

Linguists use similar structures with unequal branch lengths to study the evolution of languages (Honkola et al., 2013; Atkinson and Gray, 2005). In computing, the size of a search tree node corresponds to the probability of it being visited.

In this general context, a useful index should be robust, universal, and interpretable. A loose definition of robustness is that small changes to the tree have only small effects on the index value, except where sensitivity is desirable; universal means that the index is defined for all rooted trees; and interpretable implies a simple, consistent interpretation, enabling meaningful comparison of any pair of rooted trees. Lemant et al. (2022) provides more rigorous, axiomatic definitions. We follow these axiomatic definitions and call a tree index with all three properties an RUI index. In practical terms, robustness implies that an index is relatively insensitive to the effects of issues such as sampling error, inferential error, omission of rare types, imperfect genetic sequencing, and incomplete resolution of ancestral relationships. All our indices are dimensionless but the diversity indices can be re-scaled in terms of the branch length unit where desired.

We begin by recalling the family of diversity indices attributed to Hill (1973). These Hill numbers are functions of a set of proportions *P* = *{p*_1_, …, *p*_*n*_*}* with 0 ⩽ *p* ⩽ 1 for all *p* ∈ *P* and 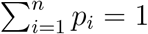 Every Hill number of order *q* ⩾ 0 can be written as

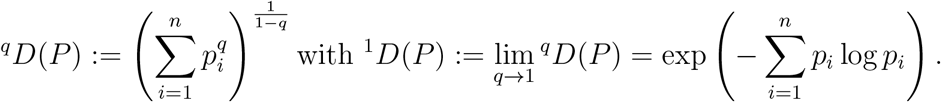

Hence ^*q*^*D* is the exponential of the Rényi entropy of order *q* (Rényi, 1961), which we will denote ^*q*^*H*, and ^1^*H* is Shannon’s entropy (Shannon, 1948). Another important special case is

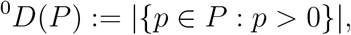

which is simply the number of types, or richness. Following Pielou (1966) and Jost (2010), we further define the evenness indices

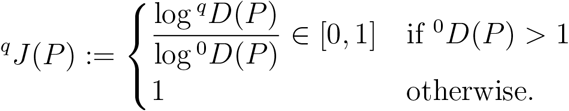

For completeness, we set ^*q*^*D*(∅) = 0 and ^*q*^*J*(∅) = 1.

We can apply these indices to a rooted tree *T* simply by equating *P* (*T*) = *{p*_1_, …, *p*_*n*_*}* to the proportional sizes of the *n* nodes of *T*, including the internal nodes. Assigning non-zero sizes to internal nodes makes sense, for example, in the case of a tumour clone tree (Noble et al., 2022). The richness index ^0^*D*(*T*) = ^0^*D*(*P* (*T*)) then quantifies the number of non-zero-sized nodes in the tree, which we will refer to as the counted nodes. In an evolutionary tree, counted nodes correspond to extant types. For each *q >* 0, the diversity index ^*q*^*D*(*T*) = ^*q*^*D*(*P* (*T*)) can be interpreted as an effective number of counted nodes, while ^*q*^*J*(*T*) = ^*q*^*J*(*P* (*T*)) gauges the evenness of the counted node sizes.

Clearly ^*q*^*D* and ^*q*^*J* are insensitive to small changes to proportional node sizes. For *q >* 0, ^*q*^*D* is also generally robust to the addition or removal of relatively small nodes (and the degree of robustness increases with *q*), whereas ^0^*D* and ^*q*^*J* are not, as is appropriate for indices that are meant to quantify richness and evenness. ^*q*^*D* and ^*q*^*J* are universal because they can be applied to any set of node sizes, and they are interpretable as described above. Yet although these indices are RUI, they are inadequate for assessing tree shape because they depend only on node sizes, ignoring both tree topology and branch lengths.

Many indices that capture aspects of tree shape have previously been defined (surveys include Pavoine and Bonsall (2011); Tucker et al. (2017); Fischer et al. (2023)) but, to the best of our knowledge, none is RUI (Table 1). We address this deficiency by developing new RUI tree indices that extend the basic indices ^*q*^*D* and ^*q*^*J* to account for tree topology and branch lengths. We do this using three types of weighted mean, which we refer to as the longitudinal mean, the node-wise mean, and the star mean (Table 2). Our consistent definitions ensure that our indices can be precisely related to each other and to ^*q*^*D* and ^*q*^*J* in numerous meaningful ways, so that all the indices belong to a single coherent system.

**Table 1.** Properties of some previously defined non-RUI tree indices (see main text for definitions and citations.)

|  | Robust |  | Universal | Interpretable |  |
| --- | --- | --- | --- | --- | --- |
|  | Robustly accounts for node sizes? | Robustly accounts for branch lengths? | Defined for all rooted trees? | Has a simple, consistent interpretation? | Can meaningfully compare any pair of rooted trees? |
| Faith's $PD$ | No | Yes | | | |
| Allen et al's $H_P$<br>Chao et al's ${}^q\bar{D}$ | Yes | | | Only for leafy ultrametric trees | |
| Sackin's index<br>Colless's index<br>Total cophenetic index | No |  |  | Yes | No |
| Lemant et al's $J^q$ | Yes | No | Yes | Only if uniform branch lengths | |

**Table 2.** Nature, notation and interpretation of RUI tree indices, including prior indices (top row) and new indices (second, third and fourth rows). Counted nodes are those with non-zero size.

| Branches or nodes | Type of average | Richness | Diversity (with $q > 0$) | Evenness (with $q > 0$) |
| --- | --- | --- | --- | --- |
| Nodes | None | ${}^0D$ = number of counted nodes | ${}^qD$ = effective number of counted nodes | ${}^qJ$ = evenness of counted node sizes |
| Branches | Longitudinal mean | ${}^0D_L$ = average branch count across the tree | ${}^qD_L$ = effective number of maximally distant leaves | ${}^qJ_L$ = evenness of branch sizes across the tree (tree symmetry if leafy and ultrametric) |
| Branches | Node-wise mean | ${}^0D_N$ = average effective outdegree, ignoring branch sizes | ${}^qD_N$ = average effective outdegree, accounting for branch sizes | ${}^qJ_N$ = tree balance |
| Branches | Star mean | ${}^0D_S$ = effective number of non-root nodes | ${}^qD_S$ = effective number of branches, accounting for branch sizes | ${}^qJ_S$ = evenness of all branch sizes |

### Further preliminary definitions

In a rooted tree, the depth of a node is the sum of the branch lengths along the unidirectional path from the root to the node. The height of the tree is the maximum depth of its non-zero-sized nodes. Nodes with no descendants are called leaves and non-leaves are called internal nodes. We define the size of a branch as the sum of the proportional node sizes that descend (directly or indirectly) from the branch. For example, in the three-leaf tree depicted in Figure 1a, the branches descending from the root have sizes 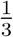 and 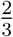, and the other two branches each have size 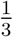. The size of any segment of a branch is the same as the size of the branch.

**Fig 1.**
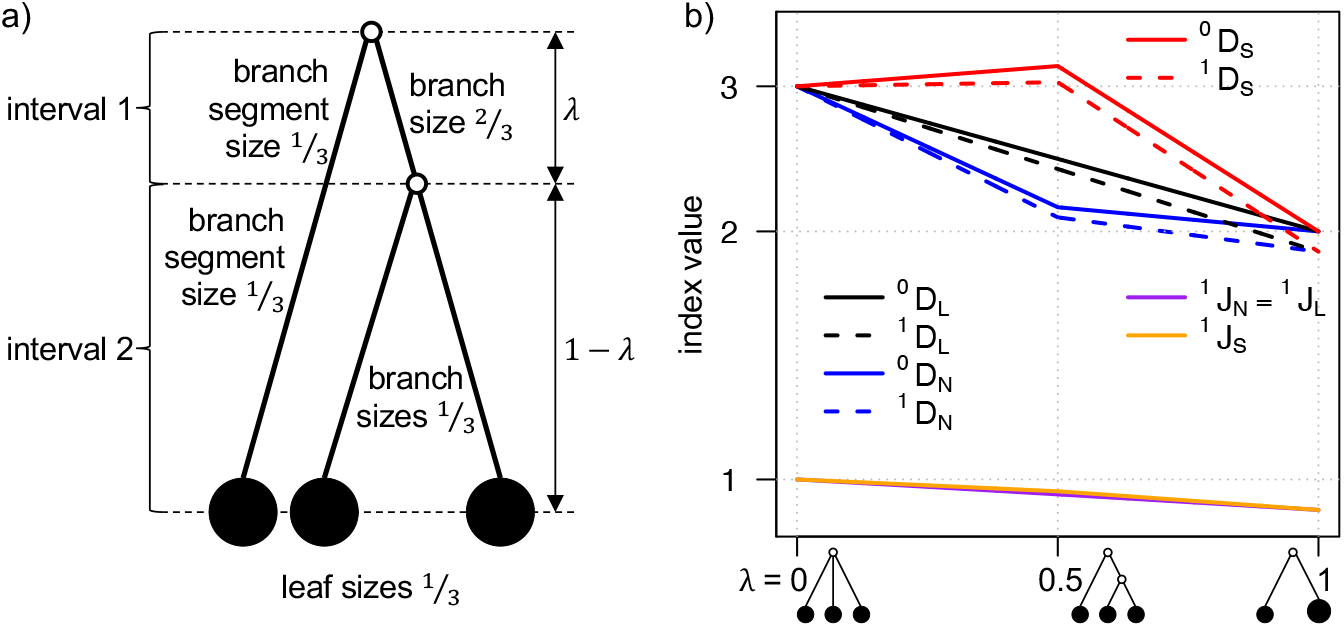
a) A leafy bifurcating ultrametric tree with three equally sized leaves. In this and every subsequent tree diagram, open circles indicate zero-sized nodes. b) Index values versus branch length *λ* for the three-leaf tree. The y-axis is log-transformed so that the curves for all diversity indices appear piecewise linear. ^1^*J*_*S*_ is slightly greater than ^1^*J*_*N*_ whenever 0 *< λ <* 1.

A leafy tree is such that all internal nodes have zero size (equivalently, all counted nodes are leaves). A tree is ultrametric if all its leaves have the same depth after the removal of all subtrees that contain only zero-sized branches (corresponding to extinct lineages in an evolutionary tree). A caterpillar tree is a bifurcating tree in which every internal node except one has exactly one child leaf. A star tree is a tree in which all non-zero-sized branches are attached to the root. We define a piecewise star tree as a tree that can be divided into transverse intervals such that, within each interval, all the non-zero-sized branches are attached to a common node. For example, the leafy ultrametric tree in Figure 1a is a star tree if *λ* = 0 or *λ* = 1 and is otherwise a caterpillar tree. To simplify our notation, we will usually omit the tree as a function argument (for example, writing ^0^*D* instead of ^0^*D*(*T*)).

It will be helpful to recall that, for a sequence of positive real numbers *X* = *x*_1_, …, *x*_*n*_, real number *r* ≠ 0, and set of positive weights *W* = *w*_1_, …, *w*_*n*_, the weighted power mean of exponent *r* is

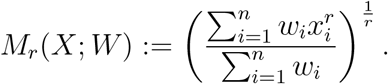

*M*_0_ is defined from the limit as

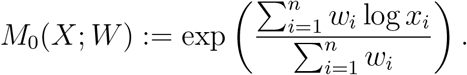

*M*_−1_, *M*_0_ and *M*_1_ are respectively the weighted harmonic, geometric, and arithmetic means. *M*_−∞_ and *M*_∞_ respectively return the minimum and the maximum. Power means are closely related to Hill numbers as, for all *q* ⩾ 0 and any sequence of proportions *P*,

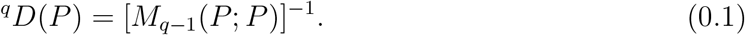

### Prior tree balance and imbalance indices

The most popular conventional tree imbalance indices can be expressed in the form

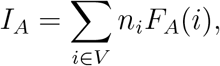

where *V* is the set of all internal nodes and *n*_*i*_ is the number of leaves that descend from node *i*, and *F*_*A*_(*i*) is a function that defines a particular index. For *I*_*S*_ (Sackin’s index), *I*_*C*_ (Colless’ index) and *I*_Φ_ (the total cophenetic index) we have

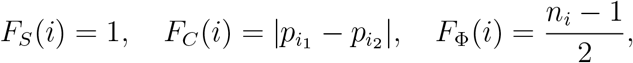

where 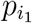 is the proportion of the *n*_*i*_ leaves that descend from the left child branch of *i*, and 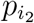 is the proportion that descend from the right child branch. Being imbalance indices, these three indices assign higher values to less balanced trees. *I*_*C*_ is defined only for bifurcating trees (in which all internal nodes have outdegree two). *I*_*S*_ and *I*_Φ_ are defined only for trees in which all internal nodes have outdegree greater than one. By convention, each index is normalized over the set of trees on *n >* 2 leaves by subtracting its minimum value over such trees and then dividing by the difference between its maximum and its minimum. The minima of *I*_*S*_, *I*_*C*_ and *I*_Φ_ are *n*, 0 and 0, and the maxima are 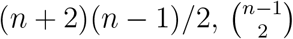 and 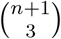, respectively (Shao and Sokal, 1990; Rogers, 1993; Mir et al., 2013).

Lemant et al. (2022) proposed instead defining tree balance or imbalance indices in the form of the weighted arithmetic mean

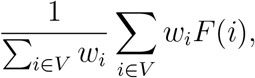

where *w*_*i*_ is the weight assigned to node *i*, and *F* (*i*) quantifies the degree to which node *i* splits its descendants into equally sized subtrees. For example, we can obtain an alternative normalization of Colless’ index by setting *w*_*i*_ = *n*_*i*_ and *F* (*i*) = *F*_*C*_(*i*). The normalizing factor 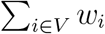 is then Sackin’s index. An advantage of this approach is that it allows us to compare the balance of any pair of trees for which *F* is defined, rather than only trees with equal leaf counts.

### Definition of the normalizing factor 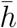

Consistent with Lemant et al. (2022), our new index definitions are based on weighted means. Our preferred weights require us to define the normalizing factor

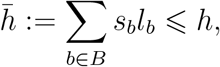

where *B* is the set of all branches in the tree, *s*_*b*_ ∈ [0, 1] is the size of branch *b, l*_*b*_ is the length of branch *b*, and *h* is the tree height. We can interpret 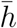 (denoted 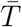 in Chao et al. (2010)) as the effective tree height or as the average counted node depth. In computer science, 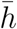 is called the weighted path length (Albers and Westbrook, 2005). For leafy trees with uniform leaf sizes and uniform branch lengths, 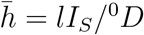, where *l* is the branch length and *I*_*S*_ is Sackin’s index. Hence 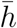 can also be considered a generalization of Sackin’s index. Indeed, we have previously argued that Sackin’s index is best interpreted not as a general imbalance index but rather as a normalizing factor, which works as an imbalance index only in the special case of trees with uniform node sizes, uniform branch lengths, and uniform outdegree (Lemant et al., 2022). 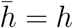 if and only if the tree is leafy and ultrametric.

### Definition of the longitudinal mean

The basic idea of the longitudinal mean is that we split the tree into transverse intervals, calculate an index value based on the proportional sizes of the branch segments within each interval, and then take a weighted average of these within-interval index values. Let *I* denote the set of transverse intervals created by locating an interval boundary at every node depth (dashed lines in Figure 1a), excluding intervals that contain only zero-sized branches. Each interval *i* ∈ *I* then contains a set *B*_*i*_ of branch segments, all of the same length, which we will refer to as the interval height *h*_*i*_. Let

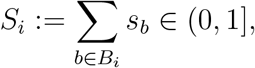

where *s*_*b*_ is the size of branch segment *b*. Then *S*_*i*_ = 1 for all intervals *i* if and only if the tree is leafy and ultrametric. It follows that

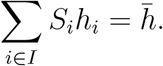

Now for each *b* ∈ *B*_*i*_, define the within-interval proportional branch size *p*_*b*_ := *s*_*b*_*/S*_*i*_ and let *P*_*i*_ := *{p*_*b*_ : *b* ∈ *B*_*i*_, *p*_*b*_ *>* 0*}*. Then 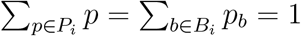 for all intervals *i* ∈ *I*.

Finally, for index *F* and tree *T*, we define the longitudinal mean of order *r* of *F* as the functional *F* ↦ *M*_*long,r*_(*F*) such that

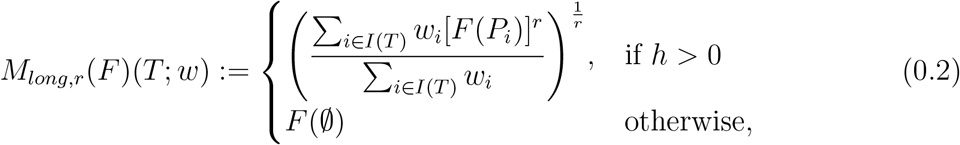

where the weight *w >* 0 is a function of *i* that remains to be specified. Hence *M*_*long,r*_(*F*) is a weighted power mean of the *F* values assigned to the intervals. For succinctness, we will omit the argument *T* and specify *w* only where necessary.

Example 0.1 For the function 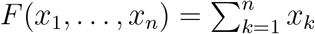 we have

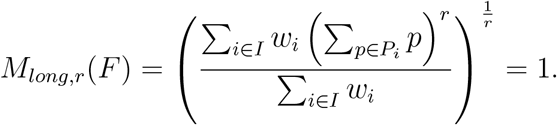

### New longitudinal mean indices

We define new tree indices as longitudinal means of ^*q*^*D* and ^*q*^*J* with *w*_*i*_ = *S*_*i*_*h*_*i*_, so that the index value assigned to each interval *i* is weighted by the product of the length *h*_*i*_ and the summed sizes *S*_*i*_ of the branch segments that *i* contains. First, we define

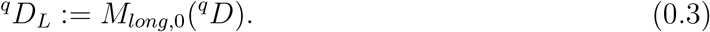

This is equivalent to ^*q*^*H*_*L*_ = *M*_*long*,1_(^*q*^*H*) with *D*_*L*_ = exp *H*_*L*_. In particular,

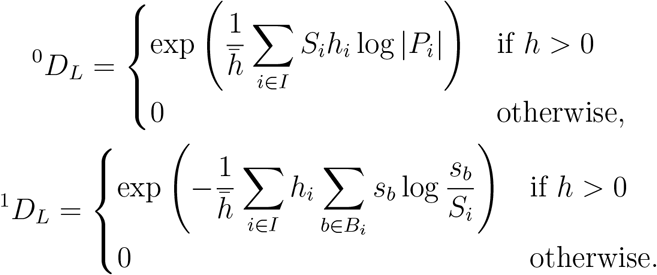

We can interpret ^0^*D*_*L*_ as the average tree width or, more precisely, as the geometric mean number of branches counted across the tree. In an evolutionary tree where branch lengths correspond to elapsed time, ^0^*D*_*L*_ equates to average richness across time, excluding extinct lineages. For *q >* 0, ^*q*^*D*_*L*_ can be interpreted as the effective number of counted nodes maximally distant from the root or – because all maximally distant counted nodes must be leaves – as the effective number of maximally distant leaves. In biological terms, this corresponds to the effective number of extant types maximally distinct from the root type.

Second, we define

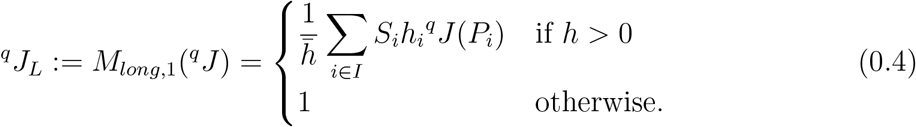

Just as ^*q*^*J* measures the evenness of node sizes, so ^*q*^*J*_*L*_ measures the average evenness of branch sizes across the tree. If the tree is leafy and ultrametric then ^*q*^*J*_*L*_ = 1 for *q >* 0 if and only if the tree is fully symmetric. Hence, when applied to leafy ultrametric trees, ^*q*^*J*_*L*_ can be interpreted as a symmetry index (also known as a sound balance index (Mir et al., 2018)).

Figure 1b illustrates how ^0^*D*_*L*_,^1^ *D*_*L*_ and ^1^*J*_*L*_ (and other index values yet to be defined) vary with branch length *λ* for the three-leaf tree of Figure 1a.

### Definition of the node-wise mean: first special case

In the special case in which all branches have the same length *l*, we can obtain a node-wise mean by calculating an index value for each node, based on the node’s child branch sizes, and then taking a weighted average of these node index values. We previously used this approach to define new tree balance indices (Lemant et al., 2022).

Let *V* denote the set of all internal nodes, excluding nodes with only zero-sized descendants. Let *C*_*i*_ denote the subtree containing only *i* and its children. For *i* ∈ *V* and *b* ∈ *C*_*i*_, let *s*_*b*_ denote the size of *b* and define

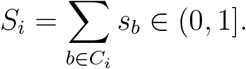

Then *S*_*i*_ = 1 for all nodes *i* if and only if the tree is a leafy piecewise star tree. It follows that

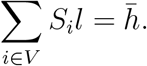

Now for each *b* ∈ *C*_*i*_, define the proportional branch size *p*_*b*_ := *s*_*b*_*/S*_*i*_ and let

*P*_*i*_ := *{p*_*b*_ : *b* ∈ *C*_*i*_, *p*_*b*_ *>* 0*}*. We then define the node-wise mean of order *r* of index *F* as the weighted power mean of the *F* values assigned to the nodes:

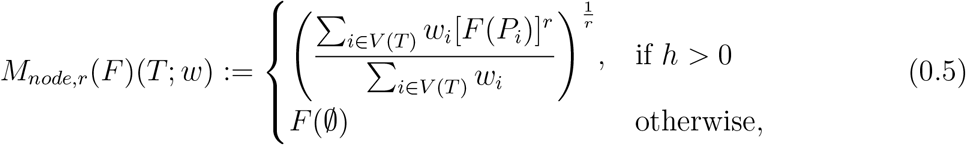

where the weight *w >* 0 is a function of *i* that remains to be specified.

### Definition of the node-wise mean: second special case

In the case of a piecewise star tree with *h >* 0, we can set the index value of each internal node *k* as the longitudinal mean index value of the subtree *C*_*k*_. We then have

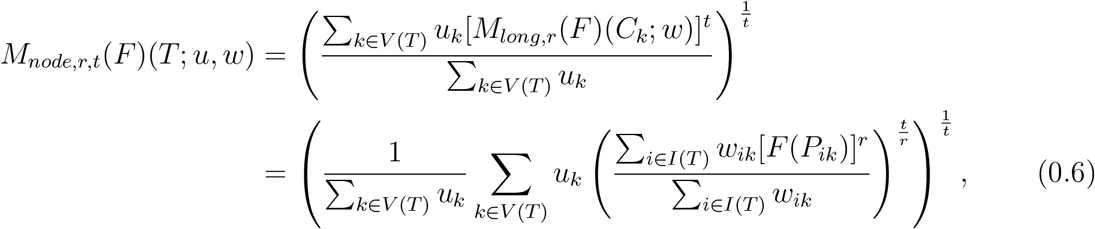

where *t* is the exponent of the across-nodes power mean, *u*_*k*_ *>* 0 is the weight assigned to node *k, P*_*ik*_ contains the proportional sizes of all branch segments that belong to both subtree *C*_*k*_ and interval *i*, and *w*_*ik*_ *>* 0 is the weight assigned to *k* associated with interval *i*.

To keep our system internally consistent we would like, in the case of piecewise star trees, the node-wise mean of any index to be equal to the longitudinal mean of the same index. Comparing Equation 0.6 with the definition of the longitudinal mean (Equation 0.2), we see that the right-hand sides are equivalent if and only if three conditions hold:

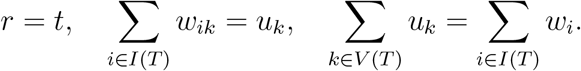

Under these conditions, summing index values across subtree intervals and then across nodes gives the same result as summing across tree intervals. We then have for any piecewise star tree *T* with *h >* 0,

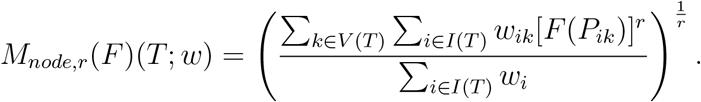

In the particular case *F* = ^*q*^*D*, the index value assigned to each node *k* (that is, the longitudinal mean index value of the subtree *C*_*k*_) measures the diversity of the child branches of *k*. When *C*_*k*_ has *m* branches of equal length and size, the node diversity of *k* is *m*. In the case *m >* 1, as one branch length is reduced towards zero while all else is kept constant, the node diversity of *k* decreases continuously to *m* − 1. Decreasing instead the size of one branch has the same effect provided *q >* 0. Hence the diversity value assigned to each node can be interpreted as an effective outdegree, and the node-wise mean diversity can be interpreted as an average effective outdegree. When *q* = 0 the effective outdegree ignores branch sizes. As *q* increases, the effective outdegree gives less weight to branches of smaller size. We would like to retain this interpretation as we generalize the definition of the node-wise mean.

### Definition of the node-wise mean: general case

In extending the definition to all rooted trees, we want to ensure that, as with the longitudinal mean, the node-wise mean changes continuously as we vary branch lengths. We illustrate this general issue with an example.

Example 0.2 Consider a leafy ultrametric tree with six leaves such that the root has two descendant branches each of length *λ*, and both non-root internal nodes have three descendant branches, all of length 1 − *λ*. When 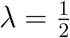 (Figure 2a), it follows from our special-case definition (Equation 0.5) that the root has richness 2, the internal nodes each have richness 3, and the node-wise mean richness is intermediate between 2 and 3. As *λ* increases from 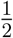 to 1, the node richness values should remain unchanged but the root node richness should be given greater weight, so that the node-wise mean richness (which we will denote ^0^*D*_*N*_) approaches 2 continuously as *λ* → 1 (Figure 2b).

**Fig 2.**
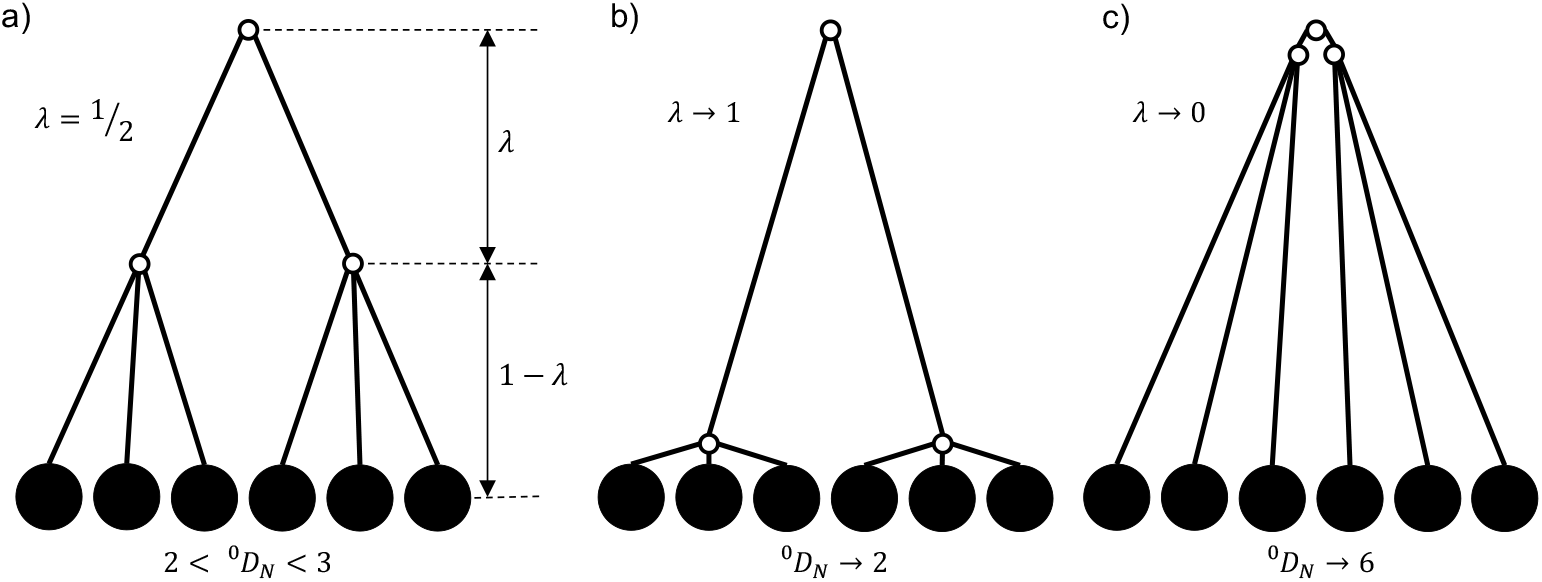
a) The six-leaf tree considered in Example 0.2 with branch length 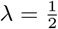. b) As *λ* → 1, the tree approaches a two-leaf star tree. c) As *λ* → 0, the tree approaches a six-leaf star tree.

At the other extreme, as *λ* decreases from 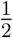 to 0, we would like ^0^*D*_*N*_ to increase continuously to 6 (Figure 2c). Given that the weight assigned to the root node richness should decrease as *λ* decreases, the only way to achieve the required increase in ^0^*D*_*N*_ is to increase the richness value assigned to each non-root internal node *k*. We can do this by making the richness value assigned to *k* depend not only on the child branches of *k* but also, to an increasing degree as *λ* decreases, on the other branches that run alongside the branches of *k*.

Generalizing from the example we conclude that, when the distance between node *k* and any ancestor *j* of *k* (in the example, the root) is less than the height of *C*_*k*_ (in the example, when 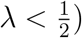), the index value assigned to *k* should depend not only on the branches of *C*_*k*_ (the child branches of *k*) but also on branch segments that descend from *j* and that coexist in transverse intervals with the branches of *C*_*k*_. The weight assigned to *k* depends only on *C*_*k*_ but the index value assigned to *k* is a weighted average of index values across *k* and all ancestors of *k*.

To formalize this concept, we first define, for interval *i* ∈ *I* and node *j* ∈ *V*,

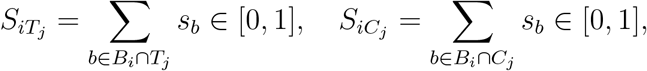

where *T*_*j*_ is the subtree containing *j* and all its descendants. This implies

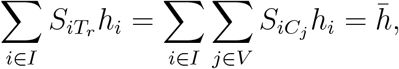

where *r* is the root (and hence *T*_*r*_ is the entire tree). 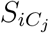 is a generalization of the *S*_*i*_ used in our previous definitions, whereas 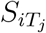 is a new concept. For each *b* ∈ *B*_*i*_ ∩ *T*_*j*_, let

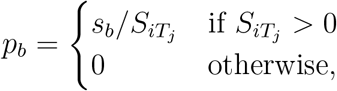

and define *P*_*ij*_ = *{p*_*b*_ : *b* ∈ *B*_*i*_ ∩ *T*_*j*_, *p*_*b*_ *>* 0*}*. We then define the node-wise average as the triple power mean

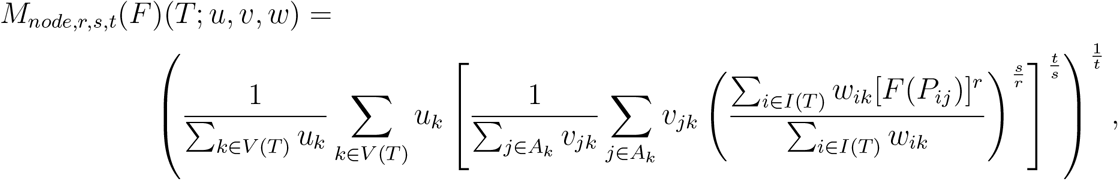

where *A*_*k*_ is the set containing *k* and all ancestors of *k, s* is the exponent of the across-ancestors power mean, and *v*_*jk*_ are the ancestor weights. This expression is consistent with Equation 0.6 if and only if

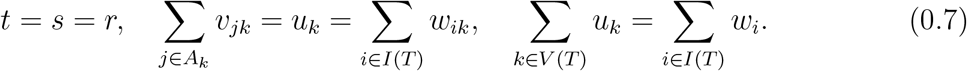

We then arrive at a simpler general definition

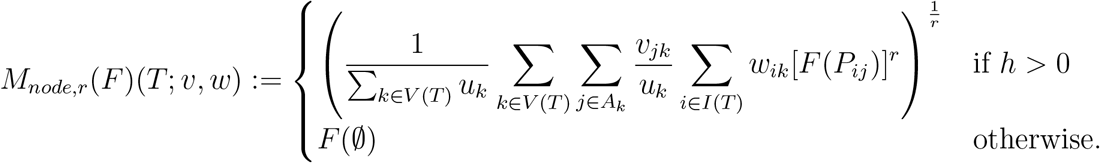

### Integral forms of the node-wise and longitudinal means

Since our preferred ancestor weights are best expressed as integrals, we will find it useful to define the longitudinal and node-wise means even more generally by integrating over depths instead of summing over intervals. Suppose we assign a non-negative density *f*_*b*_(*x*) to every branch *b* at every depth *x*, with *f*_*b*_(*x*) = 0 for every *x* at which *b* is absent. Define the tree height *h* := max*{x* : *f*_*b*_(*x*) *>* 0, *b* ∈ *B}*, where *B* is the set of all branches. We can then define branch size *s*_*b*_ as the non-increasing function of depth *x*:

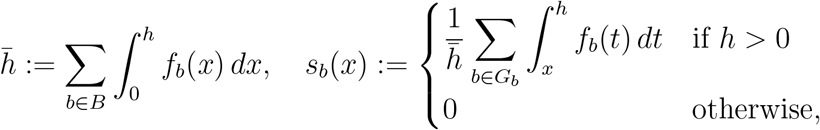

where *G*_*b*_ is the set containing *b* and all branches that descend from *b*. Let

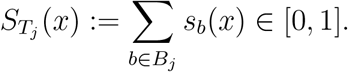

For each *b* ∈ *B*_*j*_, define the proportional branch size

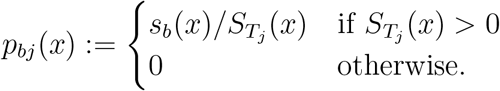

Let *P*_*j*_(*x*) := *{p*_*bj*_(*x*) : *b* ∈ *B*_*j*_, *p*_*bj*_(*x*) *>* 0*}*. We then define the node-wise mean of an index *F* as

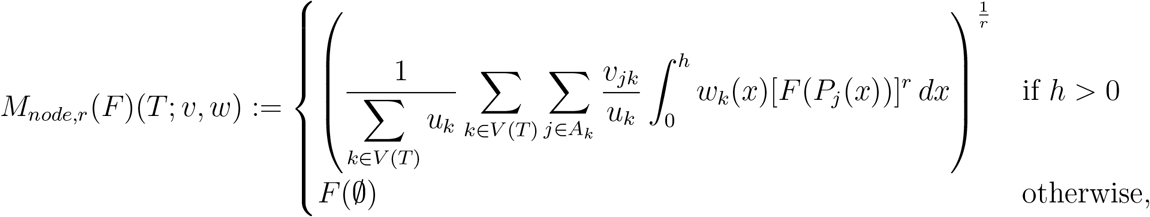

where *w*_*k*_(*x*) is the weight assigned to node *k* at depth *x*, and

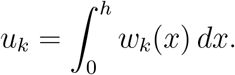

The longitudinal mean can similarly be defined in terms of integrals as

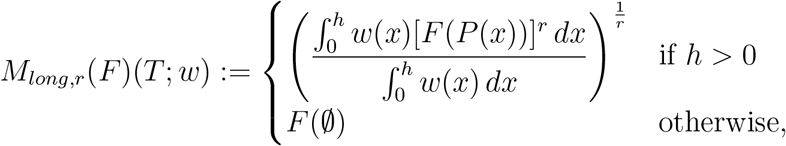

where *P* (*x*) := *{p*_*b*_(*x*) : *b* ∈ *B, p*_*b*_(*x*) *>* 0*}*,

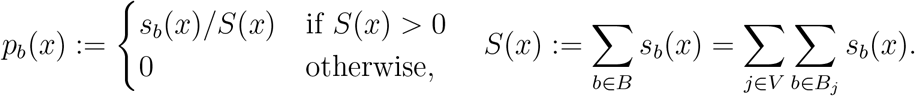

Our previous definitions are included as special cases in which the branch density is zero except at each counted node, where it is equal to the node size. In an evolutionary tree, branch density corresponds to population size, and branch size corresponds to number of extant descendants. Although it is beyond the scope of the current manuscript, we note that the integral forms would permit us to apply our indices to a more general class of tree, such that the size of any branch is allowed to vary along its length.

### New node-wise mean indices

To define new tree indices as node-wise means of ^*q*^*D* and ^*q*^*J*, we first set 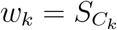,where

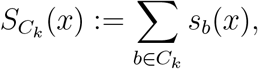

and we define the normalization factor

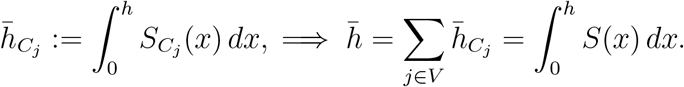

Let *d*_*k*_ denote the depth of node *k* and let *d*_*jk*_ = *d*_*k*_ − *d*_*j*_ denote the distance from *j* to *k*. Let *j*^*′*^ denote the parent of node *j*. The ancestor weight function *v* should have three properties. First, as an assumption of our general definition (Equation 0.7),

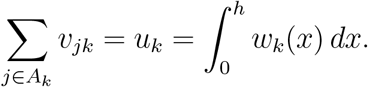

Second, *v*_*jk*_ should decrease as 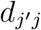 decreases. Third, *v*_*jk*_ should increase as the overlap between *C*_*j*_ and *C*_*k*_ increases. A simple way to satisfy all three conditions is to set

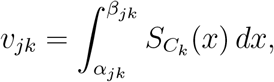

where *α*_*jk*_ := *d*_*k*_ + *d*_*jk*_ and

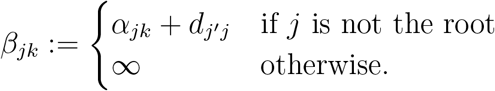

Given the above choices of *w* and *v*, we define the node-wise mean diversity of order *q* as

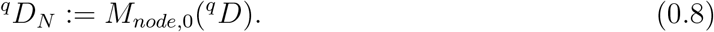

This is equivalent to ^*q*^*H*_*N*_ = *M*_*node*,1_(^*q*^*H*) with ^*q*^*D*_*N*_ = exp ^*q*^*H*_*N*_. In particular,

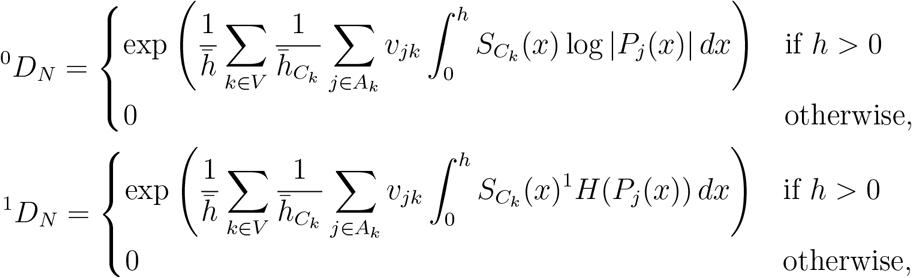

where

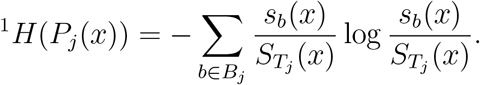

As previously explained, we can interpret ^*q*^*D*_*N*_ as an average effective outdegree (branching factor in computer science) that accounts for branch lengths only (*q* = 0) or for both branch lengths and branch sizes (*q >* 0). Less formally, ^*q*^*D*_*N*_ quantifies the bushiness of the tree.

With the same *w* and *v*, we define the universal tree balance ^*q*^*J*_*N*_ as

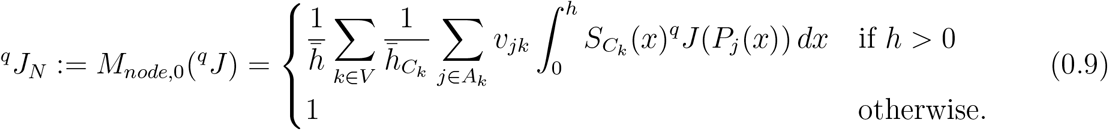

In the case of uniform branch lengths, this definition simplifies to

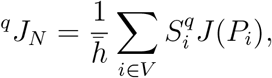

where *S*_*i*_ and *P*_*i*_ are defined as in Equation 0.5. This means that for trees with uniform branch lengths, ^*q*^*J*_*N*_ is identical to our previous definition of the tree balance index *J*^*q*^ (Lemant et al., 2022), excepting one important difference. Whereas our prior index assigns a balance score of zero to any node that has outdegree 1, the above definition instead assigns a balance score of one. Therefore linear trees are considered maximally unbalanced according to *J*^*q*^ but maximally balanced according to ^*q*^*J*_*N*_. This difference ensures that all our new evenness indices have consistent definitions and interpretations.

Example 0.3 Consider the perfectly balanced, bifurcating, leafy tree with four leaves and branch lengths *λ* (upper two branches) and 1 − *λ* (lower four branches), as shown in Figure 3a. For all *q* ⩾ 0, if 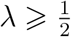 then ^*q*^*D*_*N*_ = 2, and otherwise ^*q*^*D*_*N*_ = 4^1−*λ*^, as shown in Figure 3b (dark blue curve). A step-by-step derivation is in the Appendix.

**Fig 3.**
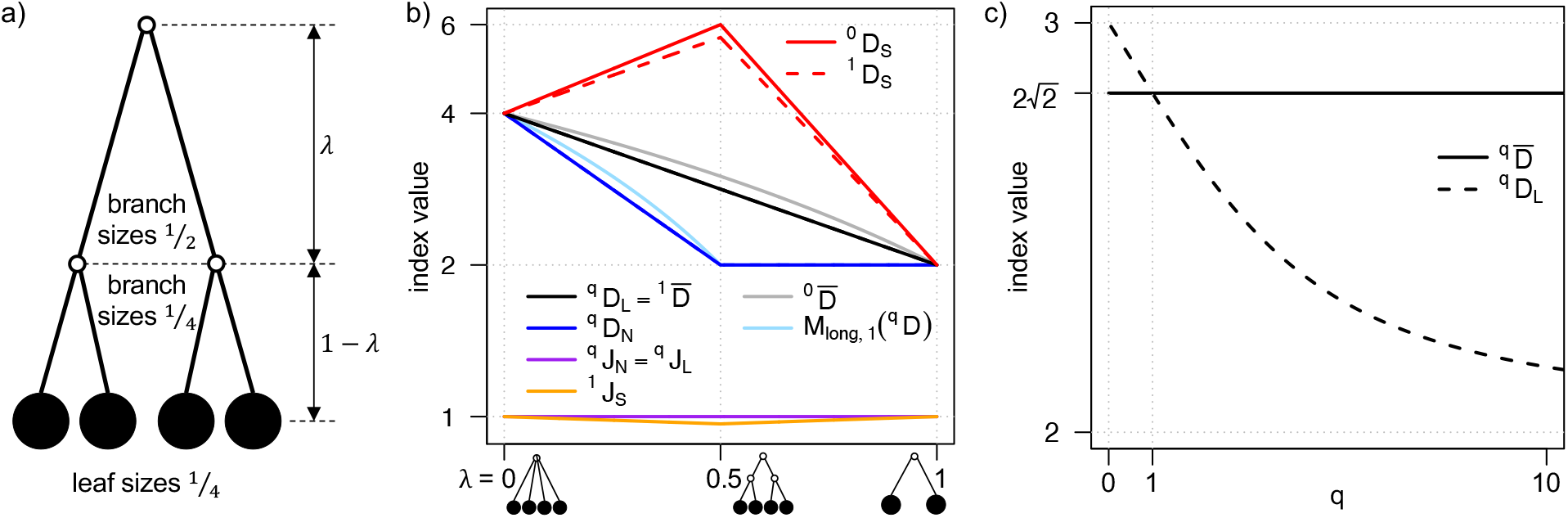
a) The four-leaf tree considered in Examples 0.3 and 0.6. b) Index values versus branch length *λ* for the tree of Example 0.6. Curves for indices with parameter *q* are independent of the value of *q* ⩾ 0. The y-axis is log-transformed so that the curves for all diversity indices except 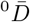 and *M*_*long*,1_(^*q*^*D*) appear piecewise linear. c) 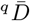 and ^*q*^*D*_*L*_ values for the four-leaf tree considered in Example 0.6, for varied *q* with 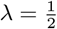.

The above example illustrates that, for leafy ultrametric trees, the node-wise mean diversity, like the longitudinal mean diversity, is a piecewise exponential function of branch lengths. Equivalently, the entropy indices are piecewise linear. This property depends on our defining the ancestor weight function *v*_*jk*_ as an integral of 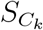. Because 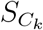 is a step function, the integrals in all our node-wise mean index definitions are simply sums of areas of rectangles, and the widths of these rectangles are linear functions of branch lengths. Our definitions are designed so that, although Equations 0.8 and 0.9 might appear complicated, in practice they produce relatively simple expressions.

### The star mean and new star mean indices

Like the longitudinal and node-wise means, the star mean is based on branch sizes. Unlike those other two means, but in common with the node-size indices ^*q*^*D* and ^*q*^*J*, the star mean ignores tree topology. The idea is that, in effect, we rearrange the tree by reattaching all branches to the root to form a star tree, while retaining branch sizes and lengths, and then calculate the longitudinal (equivalently node-wise) mean index value of the star tree. For index *F* and tree *T*, we define the star mean of order *r* of *F* such that

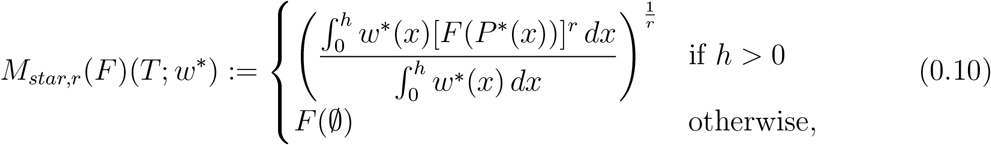

where 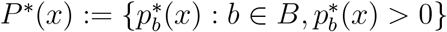,

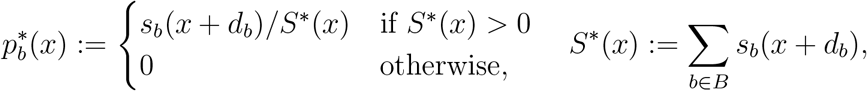

and *d*_*b*_ is the depth of the parent node of branch *b*. Note that

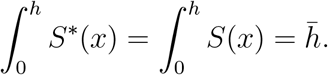

With *w*^*^ = *S*^*^, we define the star mean diversity of order *q* as

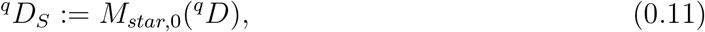

which is equivalent to ^*q*^*H*_*S*_ = *M*_*star*,1_(^*q*^*H*) with ^*q*^*D*_*S*_ = exp ^*q*^*H*_*S*_. In particular,

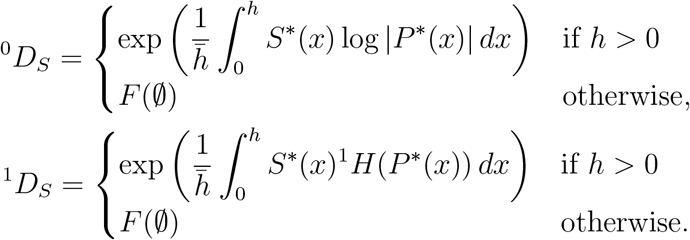

^*q*^*D*_*S*_ quantifies the effective number of branches in the tree, either accounting for branch lengths only (*q* = 0) or for both branch lengths and branch sizes (*q >* 0). Because every non-root node has exactly one parent branch, and because ^0^*D*_*S*_ accounts for branch lengths but not sizes, ^0^*D*_*S*_ can also be interpreted as an effective number of non-root nodes. We also define an index that quantifies the evenness of all branch sizes:

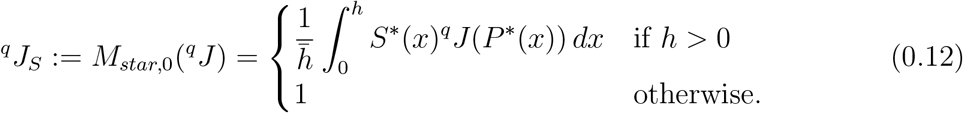

Figures 1b and 3b illustrate how ^0^*D*_*S*_,^1^ *D*_*S*_ and ^1^*J*_*S*_ values vary with branch lengths for three- and four-leaf trees.

### Non-normalized indices

Although our focus is on indices that describe shape, rather than size, we note that every longitudinal, node-wise, or star mean diversity index can be converted into a non-normalized diversity index simply by omitting the normalization factor. Such indices are useful in applications where the unit of branch length should be retained, such as when assessing loss of richness or diversity due to the removal of a node. In particular, we will find it useful to define the non-normalized entropy index

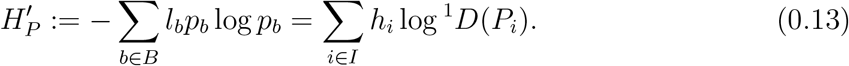

## Results

### ^q^D_L_ improves on prior indices for non-ultrametric trees

Our indices ^0^*D*_*L*_ and ^1^*D*_*L*_ are similar to well-known pre-existing indices but with important improvements (Table 3). The phylogenetic diversity of Faith (1992) – which is popular among conservation biologists – is defined as

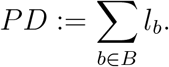

**Table 3.** Advantages of using our indices instead of previously-defined indices.

| Prior index | Proposed replacement | Equation | Advantages of replacement |
| --- | --- | --- | --- |
| Allen et al's $H_P$ | $H'_P$ | 0.13 | Interpretable for non-ultrametric trees |
| Chao et al's ${}^q\bar{D}$ | ${}^qD_L$ | 0.3 | Bounded and interpretable for non-ultrametric trees; more self-consistent; more intuitive for $q > 1$ |
| All prior tree balance and imbalance indices | ${}^qJ_N$ | 0.9 | Defined for all rooted trees; can meaningfully compare any pair of trees; accounts for node sizes and branch lengths |

Phylogenetic entropy (Allen et al., 2009) – a previous generalization of Shannon’s entropy – is defined in our notation as

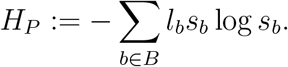

Chao et al. (2010) defined normalized versions of these indices that can be written as

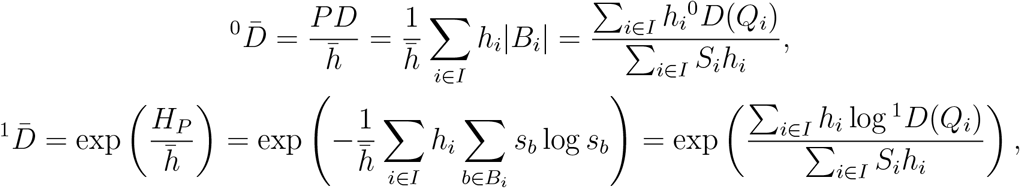

where *Q*_*i*_ = *{s*_*b*_ : *b* ∈ *B*_*i*_*}*.

A first problem with these definitions is that, for non-ultrametric trees, phylogenetic entropy lacks a clear interpretation. This issue is due to *H*_*P*_ being defined in terms of sets of branch sizes *Q*_*i*_ instead of sets of within-interval proportional branch sizes *P*_*i*_ = *{p*_*b*_ = *s*_*b*_*/S*_*i*_ : *b* ∈ *B*_*i*_*}*, as illustrated by the following example.

Example 0.4 Consider the three-node, two-leaf tree with leaf sizes *p* and 1 − *p*, and leaf depths 1 + *λ* and *λ*, respectively (Figure 4a). For this tree, as *λ* → 0,

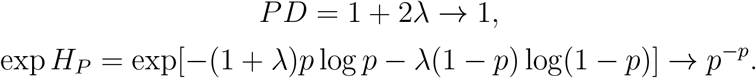

**Fig 4.**
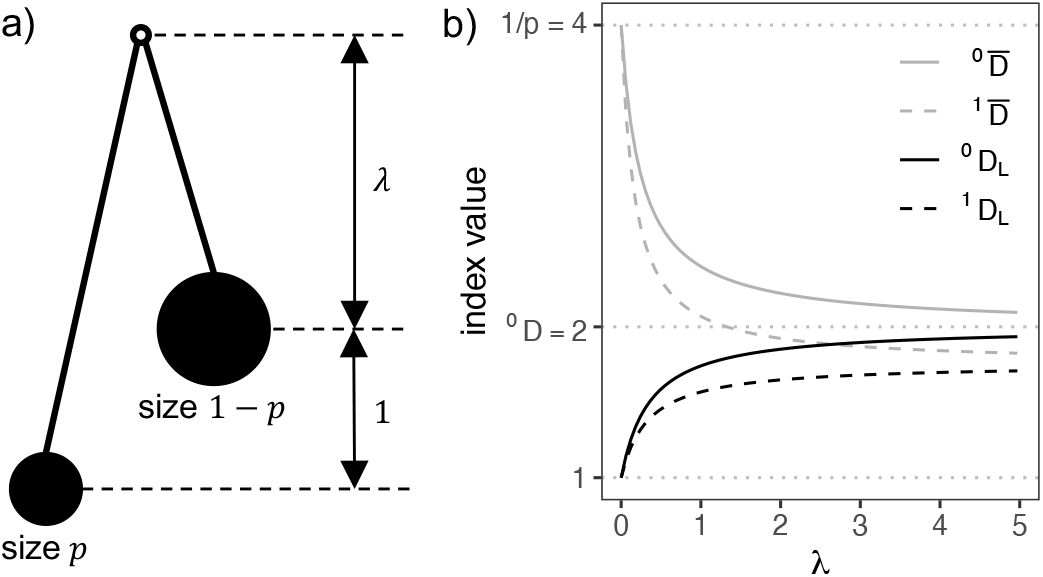
a) The two-leaf tree considered in Examples 0.4 and 0.5. b) Index values for the tree of Example 0.5 with 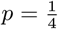. As branch length *λ* decreases, the previously defined indices 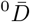 and 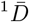 (grey curves) increase monotonically until both 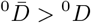 and 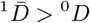. In contrast, our new indices ^0^*D*_*L*_ and ^1^*D*_*L*_ (black curves) decrease monotonically as *l* decreases, with ^0^*D*_*L*_ *<* ^0^*D* and ^1^*D*_*L*_ *<* ^0^*D* for all values of *λ*.

Therefore *PD* behaves as expected but, except when *p* = 0 or *p* = 1, exp *H*_*P*_ approaches a limit greater than 1. Hence exp *H*_*P*_ (which is supposed to be a measure of diversity) is greater than *PD* (a measure of richness). Moreover, whereas we expect diversity to be maximal when node sizes are equal, exp *H*_*P*_ is maximal when the node sizes are unequal (specifically, exp *H*_*P*_ ≈ 1.44 when *p* = *e*^−1^ ≈ 0.37). If we instead use our index 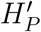 (Equation 0.13) then we obtain

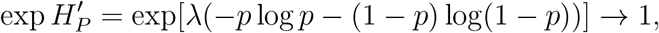

as we would expect.

A second problem is that if the tree is not ultrametric then 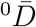 and 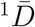 do not correspond to weighted means. If and only if the tree is leafy and ultrametric, *S*_*i*_ = 1 and *Q*_*i*_ = *P*_*i*_ for all *i* and so

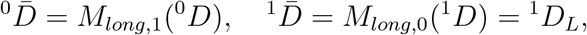

with *w*_*i*_ = *S*_*i*_*h*_*i*_ = *h*_*i*_ in both cases. Otherwise, the numerator weights *h*_*i*_ are unequal to the denominator weights *S*_*i*_*h*_*i*_. As previously noted (Chao et al., 2010; Leinster and Cobbold, 2012), this implies that 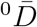 and 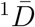 can take values exceeding the number of counted nodes when applied to non-ultrametric (or non-leafy) trees. Therefore these normalized indices lack a universal interpretation in terms of effective numbers of counted nodes (or extant types) (Leinster and Cobbold, 2012).

We avoid both problems by defining our richness and diversity indices as weighted means of the within-interval proportional branch sizes in all cases. As illustrated by the following example, the differences between ^0^*D*_*L*_ and 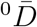 and between ^1^*D*_*L*_ and 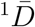 are generally unbounded and can be relatively large even when branch sizes and node sizes are not very unequal.

Example 0.5 Consider the three-node tree of Figure 4a with 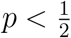. We have 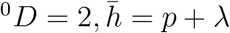, and

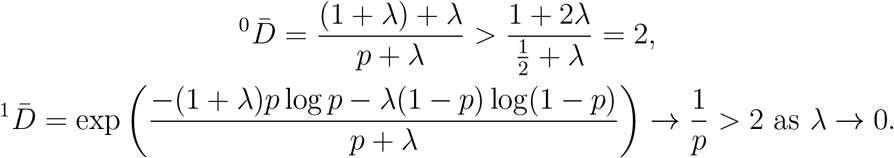

It follows that 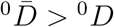 for all *λ*, and we can choose *λ* sufficiently small such that also 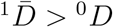 (Figure 4b, grey curves). For the same three-node tree, our new indices are instead

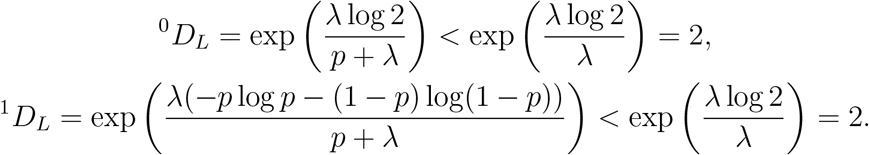

Therefore ^0^*D*_*L*_ *<* ^0^*D* and ^1^*D*_*L*_ *<* ^0^*D* for all *λ* ⩾ 0, as we would expect (Figure 4b, black curves). As *λ* → 0, both ^0^*D*_*L*_ and ^1^*D*_*L*_ approach 1, consistent with the fact that the tree has exactly one non-root node when *λ* = 0. As *λ* → ∞, the tree becomes increasingly close to being an ultrametric star tree, and hence 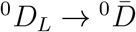 and 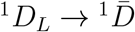 (convergence between dashed curves and between solid curves in Figure 4b).

### ^q^ D_L_ is more self-consistent and intuitive than the 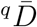 of Chao et al. (2010)

Additional problems with the 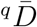 indices of Chao et al. (2010) are that they are not self-consistent, and that they have counter-intuitive properties when *q >* 1. The general definition can be expressed as

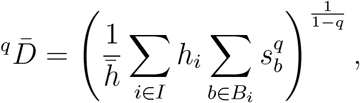

which can be restructured as

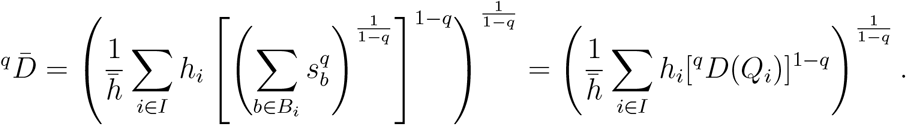

Hence for leafy ultrametric trees we have

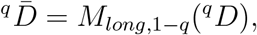

with *w*_*i*_ = *h*_*i*_ = *S*_*i*_*h*_*i*_. We have thus shown that, in the case of leafy ultrametric trees, every 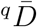 can be expressed as a weighted mean of within-interval diversities. But 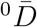 is the weighted arithmetic mean, 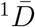 is the weighted geometric mean, and in general 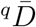 is the weighted power mean of exponent 1 − *q*. One consequence is that, for ultrametric trees in which every transverse interval contains branches of equal size, the set of within-interval values will be the same for every *q* value but the 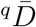 values will be different. Moreover, as *q* becomes larger, 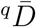 increasingly gives larger weight to *smaller* within-interval diversities. As *q* → ∞, the ^*q*^*D* value assigned to each interval approaches the reciprocal of the maximum branch size within the interval. Counter-intuitively, 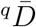 approaches the *minimum* of these within-interval ^*q*^*D* values.

These peculiar properties of 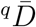 are unnecessary and have no obvious advantages.

The Hill numbers ^*q*^*D*, which are used to assign a diversity value to each interval, necessarily relate to different types of weighted mean (Equation 0.1). But the method of averaging *between* intervals need not depend on the method of calculating diversity *within* intervals. Every Hill number ^*q*^*D* can be extended to account for tree shape using the weighted arithmetic mean, the weighted geometric mean, or any other weighted power mean of the within-interval diversities by varying exponent *r* of the longitudinal mean diversity index

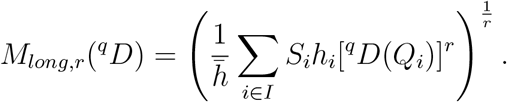

The same choice exists when defining node-wise means and star means. To avoid incompatibilities within our system, we define all our diversity indices as weighted geometric means (*r* → 0). The following example illustrates the problem and our solution.

Example 0.6 Consider again the four-leaf tree of Example 0.3 (Figure 3a). The longitudinal mean diversity values assigned to this tree are

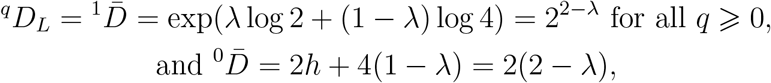

which are unequal except when *λ* = 0 or *λ* = 1 (Figure 3b, black and grey curves). In particular, in the case of uniform branch lengths 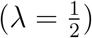, we find 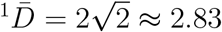 and 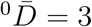 (Figure 3c, dashed curve). As derived in Example 0.3, the node-wise mean diversity for this tree is

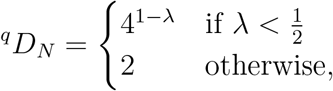

for all *q* ⩾ 0. Choosing the arithmetic mean instead of the geometric mean would instead give

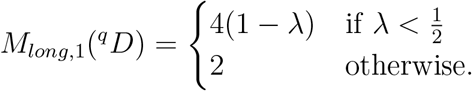

^*q*^ *D*_*N*_ := *M*_*long*,0_(^*q*^*D*)*≠ M*_*long*,1_(^*q*^*D*) for all *q* ⩾ 0 and all *λ* with 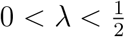 (Figure 3b, dark blue and pale blue curves). As *q* → ∞, 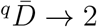 (Figure 3c, dashed curve), while ^*q*^*D*_*L*_ remains constant (Figure 3c, solid line).

### ^q^J_N_ improves on all prior tree balance and imbalance indices

As previously explained (Lemant et al., 2022) and as summarized in Tables 1 and 3, conventional tree balance and imbalance indices including Sackin’s index, Colless’ index, the total cophenetic index, and others (reviewed by Fischer et al. (2023)) have important shortcomings. In the first place, these indices account for neither node sizes nor branch lengths. This means, for example, that these indices consider all star trees maximally balanced and all caterpillar trees maximally imbalanced, even as the relative sizes of some nodes or the relative lengths of some branches approach zero (Figure 5, green lines). The tree balance index *J*^*q*^ defined by Lemant et al. (2022) varies continuously with changing node sizes but is independent of branch lengths (Figure 5, dashed purple curves). ^*q*^*J*_*N*_ improves on *J*^*q*^ by also varying continuously with branch lengths (Figure 5, solid purple curves).

**Fig 5.**
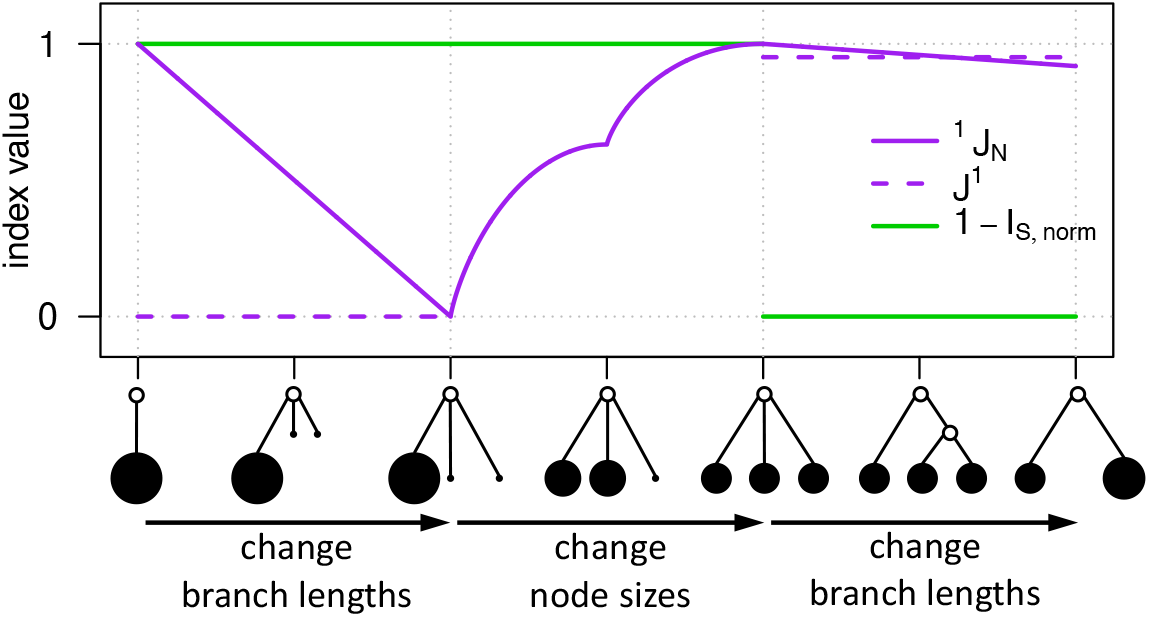
Values of three tree balance indices for a tree undergoing continuous changes. *J* ^1^ is the index introduced by Lemant et al. (2022), which is equal to ^1^*J*_*N*_ in the central third of the plot. *I*_*S,norm*_ is the normalized Sackin index, which is undefined for the leftmost, linear tree. We plot 1 − *I*_*S,norm*_ for fair comparison because *I*_*S,norm*_ is an imbalance index whereas *J* ^1^ and ^1^*J*_*N*_ are balance indices. The normalized Colless index is equal to *I*_*S,norm*_ in the rightmost third of the plot and is otherwise undefined. The normalized total cophenetic index is equal to *I*_*S,norm*_ throughout the plot.

Lemant et al. (2022) further showed that, even when restricted to the tree types on which conventional tree balance indices are defined, and even when all node sizes are equal, *J*^*q*^ enables a more meaningful comparison of trees with different degree distributions or different numbers of leaves. For example, when applied to leafy caterpillar trees with uniform branch lengths and uniform node sizes, *J*^*q*^ considers long trees (those with many leaves) to be less balanced than short ones, whereas conventional indices consider them equally imbalanced. ^*q*^*J*_*N*_, as an extension of *J*^*q*^, shares this useful property.

### Inequalities between indices

Choosing self-consistent definitions ensures that our diversity indices are related by simple sets of inequalities, which formalize and generalize the results of previous sections (Figure 6a). Hill (1973) showed that ^*q*^*D* ⩾ ^*r*^*D* for all *r* ⩾ *q* ⩾ 0. Because ^*q*^*D*_*L*_ and ^*q*^*D*_*N*_ are geometric weighted means of ^*q*^*D* values with weights independent of *q*, it follows that they obey corresponding inequalities:

Property 0.1 For all rooted trees, ^*q*^*D*_*L*_ ⩾ ^*r*^*D*_*L*_, ^*q*^*D*_*N*_ ⩾ ^*r*^*D*_*N*_ and ^*q*^*D*_*S*_ ⩾ ^*r*^*D*_*S*_ for all *r* ⩾ *q* ⩾ 0.

Additional inequalities exist between different types of diversity index but not among the evenness indices:

Proposition, 1 For all rooted trees, ^0^*D* ⩾ ^*q*^ *D*_*L*_ for all *q* ⩾ 0. For all leafy ultrametric trees, but not for all rooted trees, ^*q*^*D* ⩾ ^*q*^ *D*_*L*_ for all *q* ⩾ 0.

Proposition, 2 For all rooted trees, ^*q*^*D*_*L*_ ⩾ ^*q*^ *D*_*N*_ for all *q* ⩾ 0.

Proposition, 3 For *q >* 0, no single ordering of ^*q*^*J*, ^*q*^*J*_*L*_ and ^*q*^*J*_*N*_ applies to all leafy ultrametric trees.

**Fig 6.**
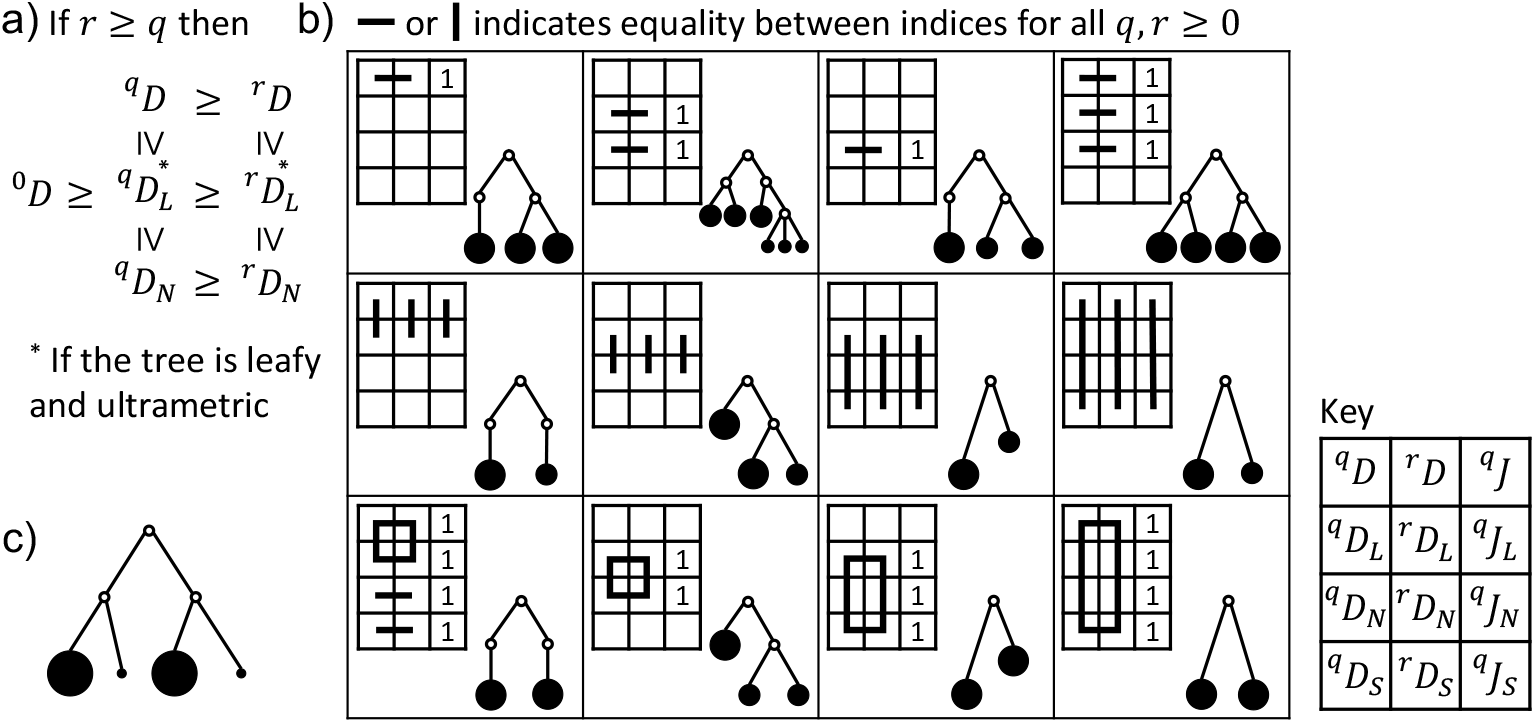
a) Inequalities between diversity indices for all *q* ⩾ 0 and all *r* ⩾ *q*. b) Examples of leafy trees with uniform branch lengths for which various index values are equal for all *q, r* ⩾ 0. The top left corner of each panel contains a grid, whose twelve squares correspond to the twelve indices shown in the key. A line connecting two grid squares indicates that the corresponding indices are equal for the tree shown in the panel. Instances where evenness indices are equal to 1 are indicated in the third grid column. c) A tree for which ^*q*^*J*_*L*_ *>* ^*q*^*J* and ^*q*^*J*_*L*_ *>* ^*q*^*J*_*N*_.

Proofs of these three propositions can be found in the Appendix. Informally, the reason why the second inequality in Proposition 1 applies only to leafy ultrametric trees is that ^*q*^*D*_*L*_, unlike ^*q*^*D*, is independent of the size of the root node (and any node arbitrarily close to the root).

### Special cases

Our consistent definitions further yield numerous simple equations that unite our indices in special cases. To simplify the statement of these results, we will assume that all branch sizes are greater than zero. This assumption implies no loss of generality because our index definitions are invariant to the addition or removal of subtrees containing only zero-sized branches (which in an evolutionary tree correspond to extinct lineages). The properties in this section hold for all *q, r* ⩾ 0.

We begin with cases in which diversities based on the same type of average but with different *q* values are equal. These first four properties, which are illustrated by simple examples in the top row of Figure 6b, follow immediately from the definitions.

Property 0.2 ^*q*^*D* = ^*r*^*D* ⇔ ^*q*^ *J* = 1 if and only if all counted nodes have equal size.

Property 0.3 ^*q*^*D*_*L*_ = ^*r*^*D*_*L*_ ⇔ ^*q*^ *J*_*L*_ = 1 if and only if the branch sizes at every depth are equal. This also implies ^*q*^*D*_*N*_ = ^*r*^*D*_*N*_ and ^*q*^*J*_*N*_ = 1.

Property 0.4 ^*q*^*D*_*N*_ = ^*r*^*D*_*N*_ ⇔ ^*q*^ *J*_*N*_ = 1 if and only if every internal node’s child branches have equal size.

Property 0.5 (^*q*^*D* = ^*r*^*D* and ^*q*^*D*_*L*_ = ^*r*^*D*_*L*_) ⇔ (^*q*^*J* = 1 and ^*q*^*J*_*N*_ = 1) if and only if the branch sizes at every depth are equal and all node sizes are equal. This implies that the tree is ultrametric and perfectly symmetric, and that ^*q*^*D*_*N*_ = ^*r*^*D*_*N*_ and ^*q*^*J*_*N*_ = 1.

In other special cases, we find equality among diversities of different types but with equal *q* values. Again, these properties are directly implied by the definitions. Simple examples are shown in the middle row of Figure 6b.

Property 0.6 ^*q*^*D*_*L*_ = ^*q*^*D* if and only if the tree is a leafy ultrametric tree in which no non-root node has outdegree greater than 1. This also implies ^*q*^*J*_*L*_ = ^*q*^*J*.

Property 0.7 ^*q*^*D*_*N*_ = ^*q*^*D*_*L*_ if and only if the tree is a piecewise star tree. This also implies ^*q*^*J*_*N*_ = ^*q*^*J*_*L*_.

Property 0.8 ^*q*^*D*_*S*_ = ^*q*^*D*_*N*_ = ^*q*^*D*_*L*_ if and only if the tree is a star tree. This also implies ^*q*^ *J*_*S*_ = ^*q*^ *J*_*N*_ = ^*q*^ *J*_*L*_.

Property 0.9 ^*q*^*D*_*S*_ = ^*q*^*D*_*N*_ = ^*q*^*D*_*L*_ = ^*q*^*D* if and only if the tree is a leafy ultrametric star tree. This also implies ^*q*^*J*_*S*_ = ^*q*^*J*_*N*_ = ^*q*^*J*_*L*_ = ^*q*^*J*.

It follows that equality both within and between types applies under more restrictive conditions, as illustrated in the bottom row of Figure 6b:

Property 0.10 ^*q*^*D*_*L*_ = ^*r*^*D* if and only if the tree is a leafy ultrametric tree with equally sized leaves in which only the root has outdegree greater than 1. This also implies ^*q*^*D*_*N*_ = ^*r*^*D*_*N*_, ^*q*^*D*_*S*_ = ^*r*^*D*_*S*_ and ^*q*^*J*_*S*_ = ^*q*^*J*_*N*_ = ^*q*^*J*_*L*_ = ^*q*^*J* = 1.

Property 0.11 ^*q*^*D*_*N*_ = ^*r*^*D*_*L*_ if and only if the tree is a piecewise star tree with equal branch sizes at every depth. This also implies ^*q*^*J*_*N*_ = ^*q*^*J*_*L*_ = 1.

Property 0.12 ^*q*^*D*_*S*_ = ^*q*^*D*_*N*_ = ^*q*^*D*_*L*_ if and only if the tree is a star tree with equally sized leaves. This also implies ^*q*^*J*_*S*_ = ^*q*^*J*_*N*_ = ^*q*^*J*_*L*_ = 1.

Property 0.13 ^*q*^*D*_*S*_ = ^*q*^*D*_*N*_ = ^*q*^*D*_*L*_ = ^*r*^*D* if and only if the tree is a leafy ultrametric star tree with equally sized leaves. This also implies ^*q*^*J*_*S*_ = ^*q*^*J*_*N*_ = ^*q*^*J*_*L*_ = ^*q*^*J* = 1.

In yet another set of special cases, the evenness formulas simplify to ratios. The following two results are immediate consequences of ^0^*D*_*L*_ or ^0^*D*_*N*_ being constant under the specified conditions.

Property 0.14 If the branch count across the tree is constant and greater than one then

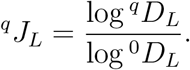

Property 0.15 If the tree has uniform outdegree greater than one and the branches present at every depth in the tree have equal lengths then

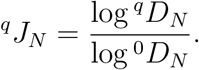

All properties described in this section would also hold if we were to define all our richness and diversity indices as weighted arithmetic, rather than geometric, means of interval or node values (or indeed any other weighted power mean). Our preference for geometric means will be justified in the next section.

### The leafy tree identity

For an important class of trees, our index definitions lead to a surprisingly simple, fundamental connection between tree balance, Shannon’s diversity index, Sackin’s index, and outdegree. This result is less obvious than the properties of the previous section and requires a more substantial proof. We term this unifying relationship the leafy tree identity.

#### Lemma 0.7

If the tree is leafy and all branches have equal length *l >* 0 then

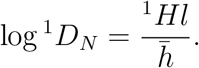

If additionally all *n* leaves have equal size then

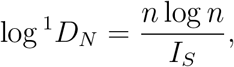

where *I*_*S*_ is Sackin’s index.

*Proof*. The proof is identical to the proof of Proposition 6 in Lemant et al. (2022), except for the base of the logarithms and the additional factor *l*.

#### Proposition 4

(The leafy tree identity; generalization of Proposition 6 in Lemant et al. (2022)) If the tree is leafy and has uniform branch lengths and all internal nodes have outdegree *m >* 1 then

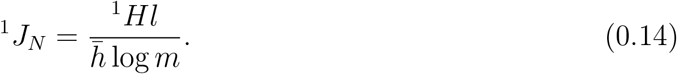

If additionally all *n* leaves have equal size then

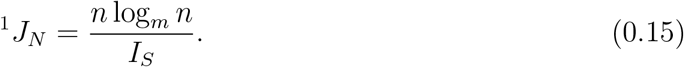

*Proof*. The result follows immediately from Lemma 0.7 and Property 0.15.

The leafy tree identity implies that, among leafy trees with uniform branch lengths and uniform outdegrees, tree balance depends only on node sizes and node depths. If two such trees have equal effective heights relative to branch length 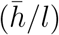, equal outdegrees (*m*), and equal node size Shannon entropy values (^1^*H*) then they must have equal balance (^1^*J*_*N*_), irrespective of topology and number of leaves. For example, Figure 7a and 7b show a pair of bifurcating leafy ultrametric trees with uniform leaf sizes and uniform branch lengths. Because these trees have equal outdegrees, leaf counts, and Sackin’s index values, the special form of the leafy tree identity (Equation 0.15) implies they must be equally balanced (other equal index values are recorded in Figure 7f). The following example applies the more general form of the leafy tree identity (Equation 0.14) to trees that are less obviously similar.

**Fig 7.**
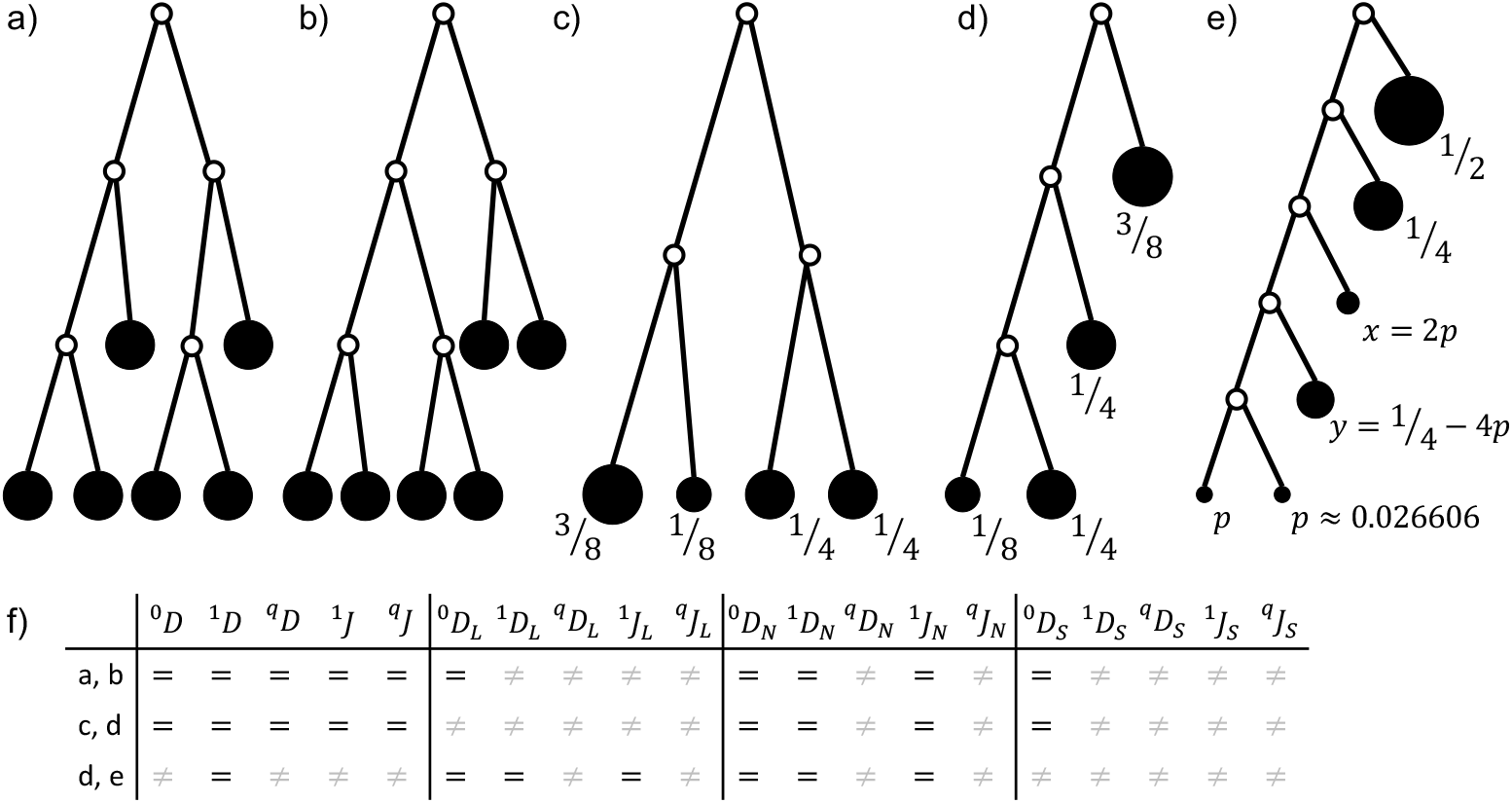
a-b) Two leafy bifurcating trees with uniform node sizes and uniform branch lengths, which differ in topology but are equally balanced. c-e) Three leafy bifurcating trees with uniform branch lengths, which differ in topology and number of leaves but are equally balanced. Nodes are labelled with their sizes. f) Table recording where pairs of trees have equal or unequal index values. Parameter *q* can take any non-negative value.

Example 0.8 Consider the bifurcating leafy ultrametric tree with four leaves, uniform branch lengths, and leaf sizes 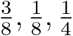 and 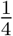 (Figure 7c). Now suppose we retain the leaf sizes but rearrange the nodes and branches to form a caterpillar tree with the node of size 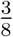 at depth *l* and one of the nodes of size 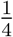 at depth 2*l* (Figure 7d). Finally, consider a six-leaf caterpillar tree with uniform branch lengths and proportional leaf sizes (in order of increasing depth) 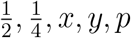 and *p*, with *p* ≈ 0.026606 (Figure 7e). All three trees have identical values of *m*, 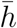 and ^1^*H* (see Appendix for derivation). Hence the leafy tree identify implies that they have equal ^1^*D*_*N*_ and ^1^*J*_*N*_ values. All three trees also have equal values of ^1^*D* = exp ^1^*H* ≈ 3.75 and ^0^*D*_*N*_ = *m* = 2. Other index values shared by pairs of trees are indicated in Figure 7f.

Equation 0.15 is especially useful because the numerator *n* log_*m*_ *n* is the minimum value that *I*_*S*_ can attain on leafy *n*-leaf trees with uniform branch lengths, uniform node sizes, and uniform outdegree *m >* 1. Hence (*n* log_*m*_ *n*)*/I*_*S*_ lies between 0 and 1 and is equal to 1 if and only if the tree is fully balanced. We previously showed (Proposition 7 in Lemant et al. (2022)) that, among all node-wise arithmetic mean indices with *w*_*i*_ = *n*_*i*_, ^1^*J*_*N*_ is the only index that satisfies Equation 0.15. Our previous proof can be straightforwardly generalized to show that Equation 0.15 cannot hold for any index of the form *M*_*node,r*_(^*q*^*J*) with *r≠* 1 or *q≠* 1. Therefore ^1^*J*_*N*_ is the only tree balance index for which this useful, unifying identity holds.

### An example cross-disciplinary application

We illustrate the universality of our methods by using them to compare the shapes of two trees from different fields of research, representing dissimilar processes and constructed using different methods. The first of these trees depicts the evolution of the Human Immunodeficiency Virus (HIV) within a host, as inferred from molecular data and as used in another recent study of tree shape indices (Barzilai and Schrago, 2023). The second tree represents the diversification of the Uralic language family (Honkola et al., 2013). To simplify the exposition we assign size zero to all internal nodes and an equal size to all leaves.

If we disregard the inferred branch lengths then it is difficult by eye to assess which tree is the more diverse or more balanced (Figure 8a, b). These apparent similarities are borne out in the shape index values (Figure 8c, d). Excepting one node, both trees are bifurcating and therefore both have ^0^*D*_*N*_ ≈ 2. The two trees have similar branch counts in total (^0^*D*_*S*_ = 33 and 32) and at each depth (3 *<* ^0^*D*_*L*_ *<* 4). The ^1^*D*_*N*_, ^1^*D*_*S*_ and ^1^*D*_*L*_ values are somewhat lower than the corresponding richness values due to imbalances, as captured by our evenness indices, which are likewise similar for the two trees (^1^*J*_*N*_ and ^1^*J*_*L*_ between 0.7 and 0.8; ^1^*J*_*S*_ ≈ 0.86). Lemma 0.7 further implies similar *I*_*S*_ values (93 and 97).

**Fig 8.**
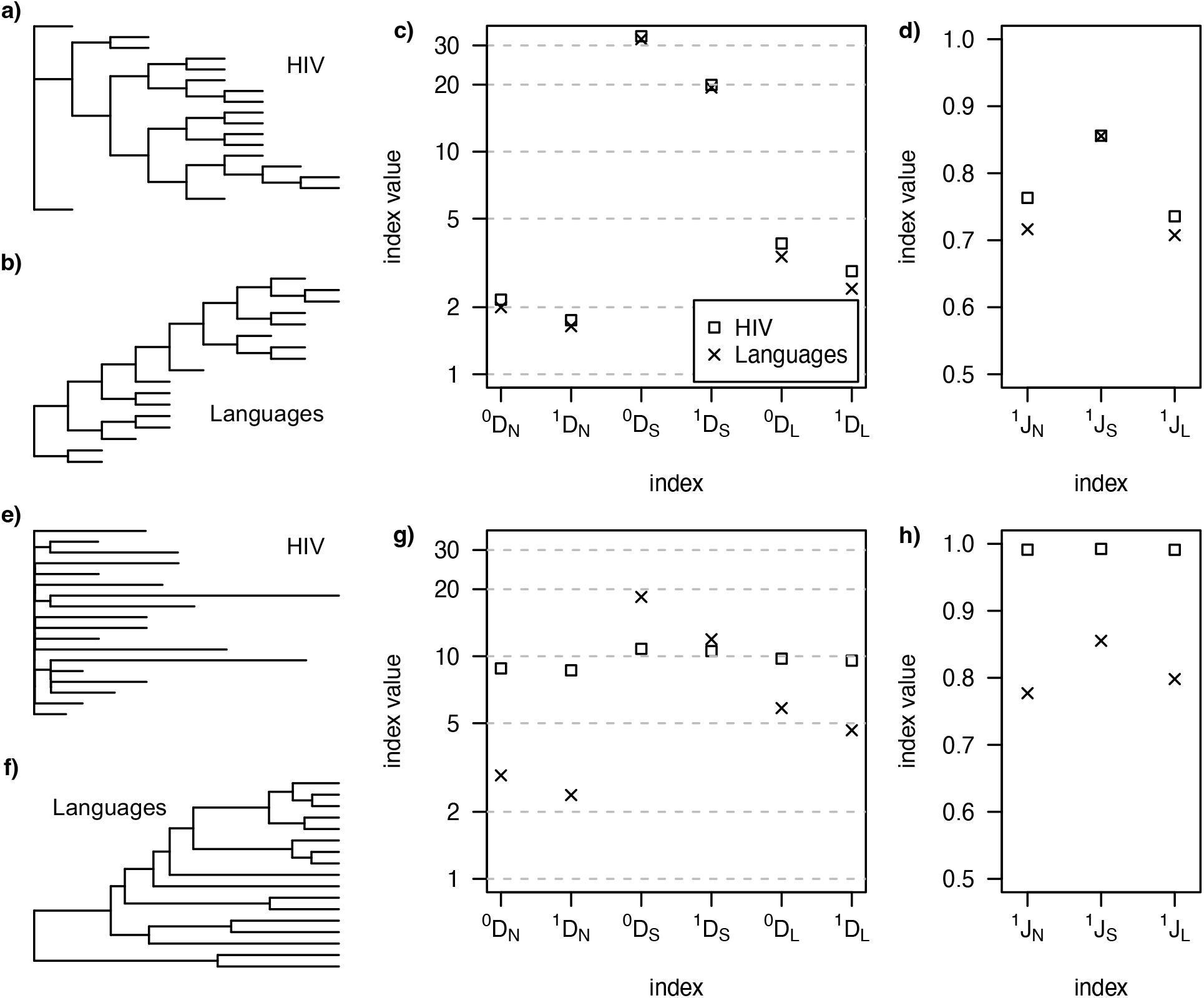
a-b) Trees with equalized branch lengths representing the within-host evolution of HIV (a) and the evolutionary history of the Uralic languages (b). c) Diversity index values for the two trees with equalized branch lengths. d) Evenness index values for the two trees with equalized branch lengths. e-f) The same trees but with the originally inferred branch lengths. g) Diversity index values, accounting for branch lengths. h) Evenness index values, accounting for branch lengths. In all cases, leaves are assigned equal size and internal nodes are assigned size zero. The HIV tree was sourced from the GitHub repository associated with Barzilai and Schrago (2023) (file PIC38051.tre) and the languages tree from the D-PLACE database (Kirby et al., 2016) (folder honkola et al2013).

When we restore the inferred branch lengths, the two trees no longer look alike (Figure 8e, f). The HIV phylogeny approximates a non-ultrametric star tree, with long branches originating close to the root. The average effective out-degree of the HIV tree, accounting for unequal branch lengths, is substantially higher than two (^0^*D*_*N*_ ≈ 9); the effective number of branches is three times lower than when branch lengths are ignored (^0^*D*_*S*_ ≈ 11); and there are more than twice as many parallel branches (^0^*D*_*L*_ ≈ 10). Because the HIV tree is approximately a star tree with equal node sizes, all its diversity indices are approximately equal and all its evenness indices are close to one (Property 0.12). In the case of the languages tree, accounting for the inferred branch lengths – which are approximately exponentially distributed and not nearly so depth-dependent – has only a small effect on most index values. The diversity indices for the languages tree remain far from equal. Altogether our indices thus show that the HIV tree is much bushier, has a larger number of effective types, and is in every sense more balanced than the languages tree (Figure 8g, h).

In summary, the clear differences between these two trees, implying different modes of evolution, are captured only by indices that account for their different branch length distributions. An analysis based on prior tree balance indices, which ignore branch lengths, would incorrectly conclude that the trees have very similar shapes and plausibly resulted from similar processes.

### An example application to model-generated trees

As a final demonstration of the potential for our indices to distinguish trees generated by different processes, we reanalyse results of a recent computational modelling study of tumour evolution by Lewinsohn et al. (2023). The original study sought to infer differences between the shapes of evolutionary trees corresponding to alternative modes of tumour expansion – boundary-driven growth (BDG) versus unrestricted growth. On average, the BDG model was found to generate ultrametric time trees with higher variance in their terminal branch lengths, and non-ultrametric gene trees with higher variance in their leaf depths (mutations per cell).

To see how our tree shape indices vary with simulated tumour growth mode, we consider the two representative simulated tumours from Figure 1 of Lewinsohn et al. (2023). The time trees (Figure 9a-c) have the same number of leaves and almost identical effective numbers of non-root nodes (^0^*D*_*S*_ ≈ 117 and 118). However, the BDG time tree has 22% higher effective branch count (^1^*D*_*S*_ ≈ 65 versus 53), 26% higher branch count across the tree (°DL ∼ 28 versus 22), and 25% higher leaf diversity (^DL ∼ 21 versus 17).

**Fig 9.**
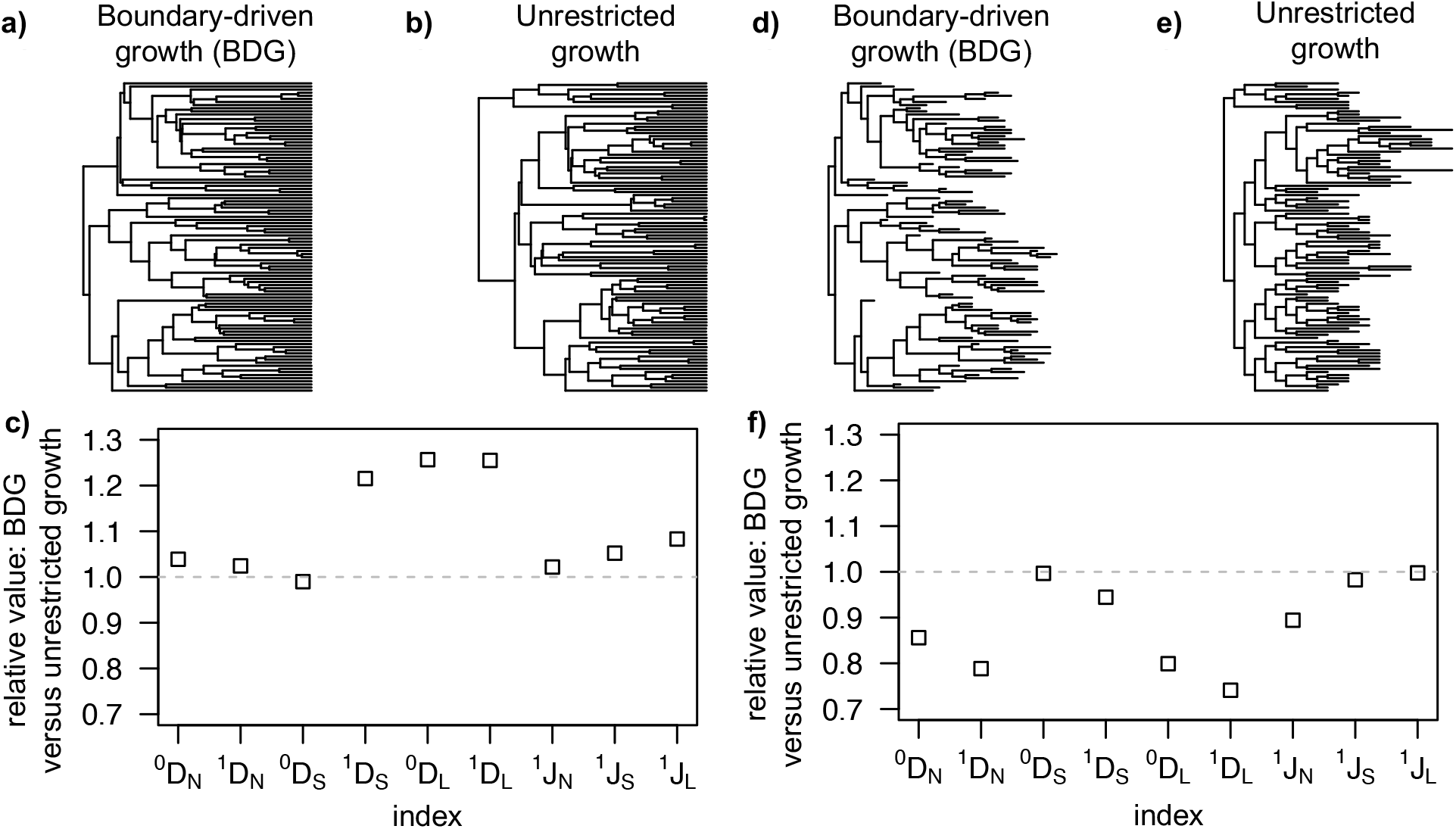
a-b) Time trees generated by computational models of tumour evolution with boundary-driven growth (a) or unrestricted growth (b). Leaves represent extant cells and branch lengths are proportional to time elapsed between cell division events, c) Tree shape index ratios for the two time trees, d-e) Gene trees generated by the same simulations as the time trees. Leaves represent extant cells and branch lengths are proportional to genetic distances, f) Tree shape index ratios for the two gene trees. All tree data was obtained from the GitHub repository associated with Lewinsohn et al. (2023).

The gene trees (Figure 9d-f) likewise have the same number of leaves and almost identical effective numbers of non-root nodes (*D*_*S*_ ≈ 136 in both cases). But the BDG gene tree, being less star-like, has substantially lower average effective outdegree (^0^*D*_*N*_ ≈ 2.6 versus 3.0; ^1^*D*_*N*_ ≈ 2.1 versus 2.6), 20% fewer branches across the tree (^0^*D*_*L*_ ≈ 17 versus 21), and 26% lower leaf diversity (^1^*D*_*L*_ ≈ 11 versus 15). The BDG gene tree is also less balanced (^1^*J*_*N*_ ≈ 0.76 versus 0.84).

Whereas well chosen problem-specific indices might give greater statistical power for distinguishing particular tree types, an advantage of our multi-dimensional system is that it is designed to be universally applicable, to facilitate comparisons between studies and data sets. Leaf depth variance, for instance, cannot by itself tell apart ultrametric trees, while terminal branch length variance is inapplicable to trees with uniform (or unknown) branch lengths.

## Discussion

The seminal paper of Hill (1973) cautions that there is “almost unlimited scope for mathematical generality in relation to measures of diversity and taxonomic difference” and therefore “Simple and well-understood indices should be used”. In accordance with this advice, here we have constructed new tree shape indices as weighted means of the most standard, basic diversity and evenness indices. This systematic approach ensures that all our indices are not only robust and universally applicable but also have simple, consistent interpretations and clear interrelationships.

Some of the indices we have defined here are refinements of prior approaches to assessing tree shape. Our ^*q*^*D*_*L*_ and 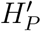 are similar to the 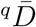 of Chao et al. (2010) and the phylogenetic entropy *H*_*P*_ of Allen et al. (2009), respectively, but are more self-consistent and can be meaningfully applied to non-ultrametric trees. ^*q*^*J*_*N*_ builds on the ideas of Lemant et al. (2022) but, by accounting for branch lengths – a key advantage of prior phylogenetic diversity and phylogenetic entropy indices, not shared by any prior tree balance indices – generalizes the concept of tree balance to a wider class of trees. These new indices share all the desirable properties but not the shortcomings of their predecessors and can therefore universally supersede them (Table 3). For the remainder of our indices describing average effective out-degree, effective numbers of nodes and branches, and evenness of branch sizes, we know of no precedents. In combination, our indices provide a more sophisticated, general, multidimensional description of tree shape than has previously been possible.

Whereas we have focussed on a system built around ^*q*^*D* and ^*q*^*J*, it is easy to use our general definitions of the longitudinal, node-wise, and star means to quantify other aspects of tree shape. A parallel, self-consistent system of indices can be defined by setting *w*_*i*_ = *h*_*i*_ instead of *w*_*i*_ = *S*_*i*_*h*_*i*_ in Equations 0.3 and 0.4, and setting *w*_*k*_ = 1 instead of 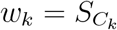 in Equations 0.8 and 0.9. These indices, which are robust to small changes in branch lengths but not node sizes, are normalized by dividing by *h* instead of 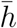 Alternatively, ^*q*^*D* can be replaced by another basic diversity index, or ^*q*^*J* by another evenness index, such as the ratio ^*q*^*D/*^0^*D* preferred by Hill (1973) (see also Smith and Wilson (1996); Jost (2010); Tuomisto (2012)). Based on the means, we can also straightforwardly derive expressions for higher moments to obtain indices that, for example, quantify how much effective out-degree varies across all nodes or varies with node depth.

There are nevertheless several reasons for preferring our specific definitions. First, the foundational ^*q*^*D* and ^*q*^*J* are the most popular diversity and evenness indices among biologists (Tucker et al., 2017; Tuomisto, 2012). Second, defining entropy and evenness indices as weighted arithmetic means, and diversity indices as weighted geometric means, results in relatively simple expressions, especially in the case of leafy ultrametric trees.

Third, ^1^*J*_*N*_ is the only universal tree balance index for which the unifying leafy tree identity holds. In summary, we have taken the best of the existing indices, improved them, unified them, and filled in the gaps to create a coherent system (Table 2).

Given the ubiquity of tree structures, we expect our multidimensional method of describing tree shape to empower research and inform decision making in diverse domains. Our initial development of universal, robust indices was motivated by the need to compare and categorize non-leafy, non-ultrametric trees representing the clonal evolution of human tumours, where node sizes (corresponding to cell subpopulation sizes) and branch lengths (genetic distances) convey valuable information (Noble et al., 2022). Tree structures with node sizes and branch lengths are likewise centrally important in community ecology, conservation biology, systematic biology, and the study of microbial evolution. For instance, our indices can be used instead of conventional tree balance indices to evaluate alternative models of speciation, or to investigate how the mode of evolution of a pathogenic virus varies with geographical location, time period, or strain. In place of phylogenetic diversity and phylogenetic entropy, our non-normalized diversity indices could be used to inform policy making by quantifying how different actions would affect biodiversity. Beyond biology, obvious subjects for analysis include phylogenetic trees of language evolution, hierarchical organizational structures, and the tree data structures that abound in computing. As we have illustrated, our generic indices can be used not only within but also across domains to uncover similarities and differences in, say, the evolution of organisms, languages, and technologies.

One key topic for further theoretical research is to derive the expected values and covariances of our indices under standard tree generation models, such as the uniform model and the Yule process, for comparison with empirical data. Relationships between our indices and distance-based metrics such as the mean pairwise distance (which lacks a universal normalization (Tsirogiannis et al., 2012)) also remain to be examined. In the same vein as Figure 7, we are investigating sets of distinct trees to which our indices assign equal values, to determine whether additional indices might ever be needed to distinguish between trees in typical applications. Towards establishing a universal standard for describing tree shape, we are developing software packages for calculating index values that can be integrated with popular tree inference methods. Just as the first step in analysing a set of measurements is to calculate the mean and variance, so we propose that, whenever one encounters a rooted tree, a useful first step will be to describe its shape by evaluating our indices.

## Acknowledgements

We are grateful to Kerry Manson for helpful comments on an earlier draft of this manuscript, to Lucia Barzilai and Chiara Barbieri for helping us obtain suitable empirical tree data, and to anonymous reviewers for suggesting various improvements.

## Data, and code availability

All data sets used in this study have previously been published; the captions of Figures 8 and 9 provide precise references. An open source R package to calculate our new tree shape indices for trees in Newick, NEXUS or phylo format is at https://github.com/kimverity/RUIindices.

## APPENDIX

### Derivation of ^q^D_N_ in Example 0.3

For the root *r*, we have 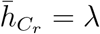 and *A*_*r*_ = *{r}*. For either of the other internal nodes *k*, we have 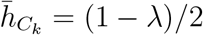 and *A*_*k*_ = *{k, r}*. The subtree weights are

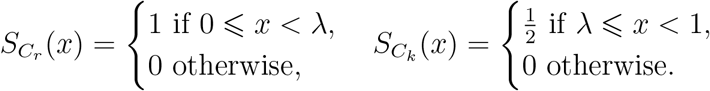

The ancestor weights are

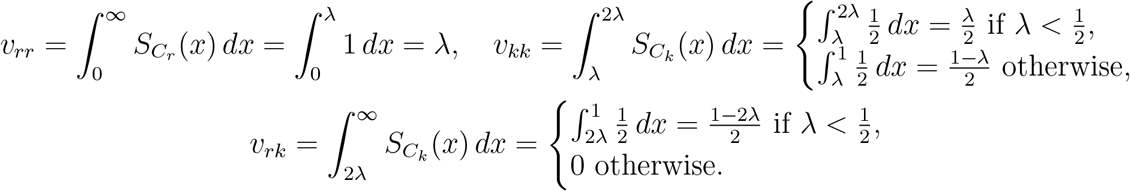

The node diversity values are, for all *q* ⩾ 0,

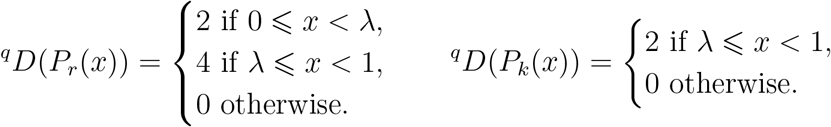

Hence

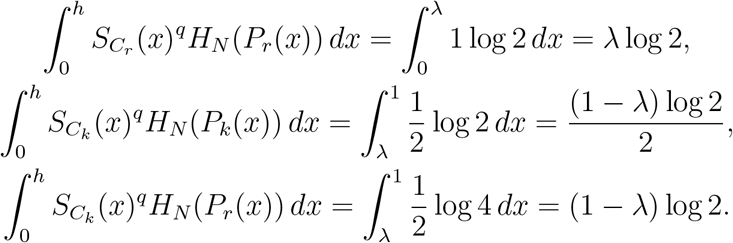

For all *q* ⩾ 0, it follows from Equation 0.8 that if 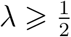 then

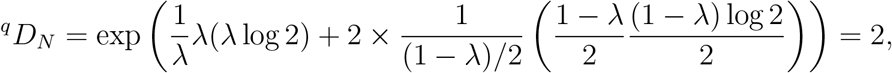

and otherwise

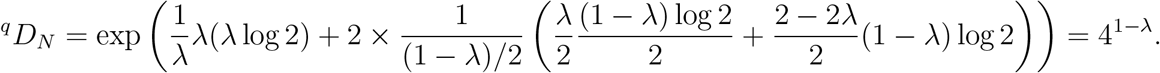

#### Proof of Proposition 1

*Proof*. For *q* ⩾ 0, let *k*(*q*) ∈ *I*(*T*) such that ^*q*^*D*(*P*_*k*(*q*)_) ⩾ ^*q*^ *D*(*P*_*i*_) for all *i* ∈ *I*(*T*), and let 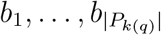 denote the non-zero-sized branches in the interval *k*(*q*). Then, by a basic property of generalized means,

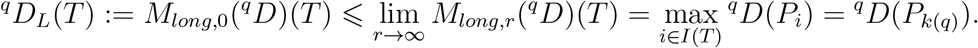

For the first part of the proposition, we note that for any interval *i* ∈ *I*(*T*), the number of non-zero-sized branches in *i* is |*P*_*i*_|, which cannot exceed the number of counted nodes ^0^*D*(*T*). Hence, for any rooted tree *T* and all *q* ⩾ 0,

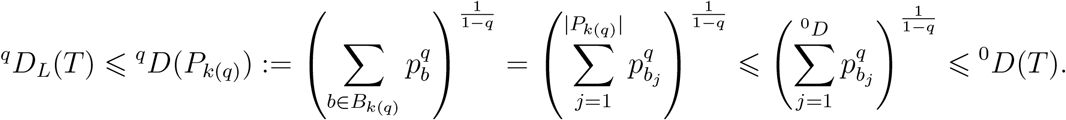

We now turn to the second part. By definition, for all *i* ∈ *I*, if *b* ∈ *B*_*i*_ then branch size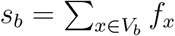, where *V*_*b*_ is the set of all nodes that descend from *b*, and *f*_*x*_ is the proportional size of node *x*. For all rooted trees we have 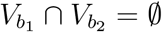 for all *b*_1_, *b*_2_ ∈ *B*_*i*_ with *b*_1_*≠ b*_2_. For ultrametric trees, 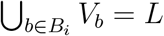, where *L* is the set of all leaves in the tree. For leafy ultrametric trees, *S*_*i*_ = 1 for all *i* and hence *p*_*b*_ = *s*_*b*_ for all *b* ∈ *B*_*i*_. Then for any leafy ultrametric tree *T* and all *q* ⩾ 0,

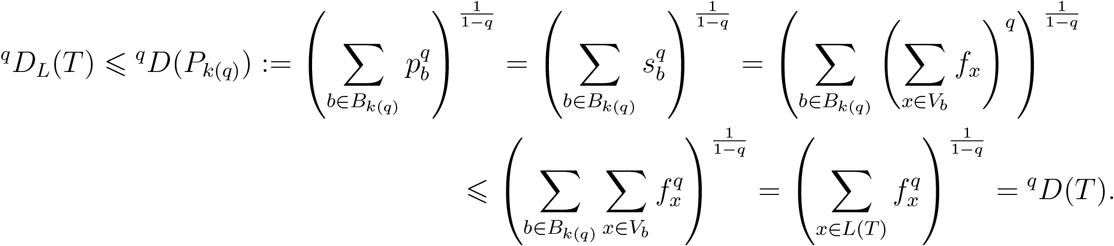

Finally we will prove that this inequality does not hold for all rooted trees. We will do so in a more general context to show that the result is independent of our choice of weight function *w* and exponent *r*. Let ^*q*^*D*_*L,r,w*_ = *M*_*long,r*_(^*q*^*D*; *w*) for real *r*, where *w* is a continuous, monotonically increasing function of *S*_*i*_ and *h*_*i*_ such that *w*_*i*_ = *w*(*S*_*i*_, *h*_*i*_) *>* 0 when *S*_*i*_ *>* 0 or *h*_*i*_ *>* 0, and *w*_*i*_ → 0 as *S*_*i*_ → 0 or *h*_*i*_ → 0. First consider the leafy but non-ultrametric three-leaf star tree *T*_1_ in which one leaf has size 1 − *p* and depth *λ*, and the other two leaves have size 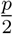 and depth 1 + *λ* (as in Figure 4 but with one more leaf). Now

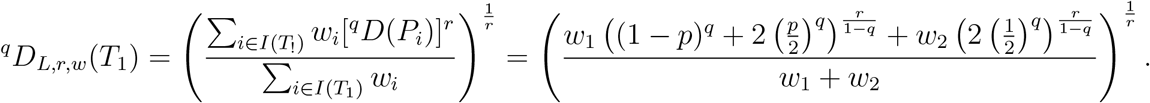

Since *w*_1_ depends only on *λ* and *w*_2_ depends only on *p*, we can make *λ* a function of *p* such that 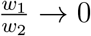 as *λ* → 0 and *p* → 0, in which case ^*q*^*D*_*L,r,w*_(*T*_1_) → 2 as *λ* → 0 and *p* → 0. Also, for all *q >* 0, ^*q*^*D*(*T*_1_) → 1 as *p* → 0. Hence ^*q*^*D*_*L,r,w*_(*T*_1_) *>* ^*q*^*D*(*T*_1_) as *λ* → 0 and *p* → 0. Instead setting *λ* = 0 makes *T*_1_ ultrametric but non-leafy, with ^*q*^*D*_*L,r,w*_(*T*_1_) → 2 and ^*q*^ *D*(*T*_1_) → 1 as *p* → 0 as before.

#### Proof of Proposition 2

*Proof*. For every node *k*, every *j* ∈ *A*_*k*_, and at every depth *x*, we have *P*_*j*_(*x*) ⊆ *P* (*x*) and so ^*q*^ *H*(*P*_*j*_(*x*)) ⩽ ^*q*^ *H*(*P* (*x*)) for all *q* ⩾ 0. Hence

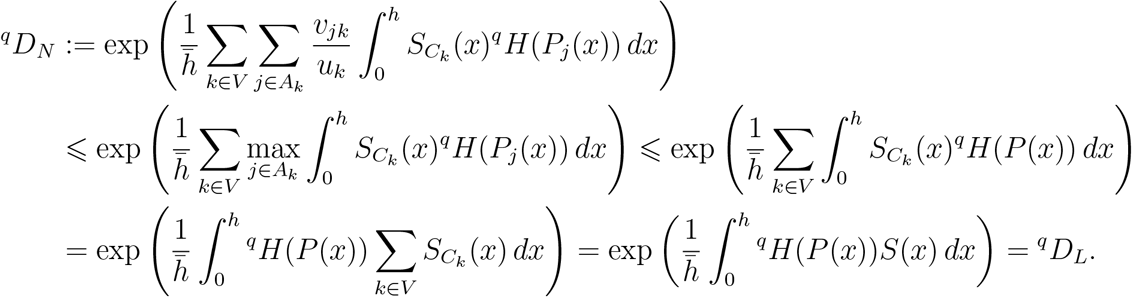

#### Proof of Proposition 3

*Proof*. The top left panel of Figure 6b shows a leafy ultrametric tree for which ^*q*^*J* = 1, ^*q*^*J*_*L*_ *<* 1 and ^*q*^*J*_*N*_ *<* 1. The third panel in the top row of Figure 6b shows a leafy ultrametric tree for which ^*q*^*J <* 1, ^*q*^*J*_*L*_ *<* 1 and ^*q*^*J*_*N*_ = 1. Now consider the four-leaf, bifurcating, leafy ultrametric tree with uniform branch lengths, such that the sizes of each pair of sibling leaves are *ϵ* and 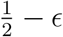 (Figure 6c). As *ϵ* → 0, we have 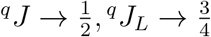 and 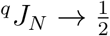. The different orderings of ^*q*^*J*, ^*q*^*J*_*L*_ and ^*q*^*J*_*N*_ for three trees are inconsistent with any universal ordering.

#### Derivation of index values in Example 0.2

Since the tree of Figure 7c is ultrametric, 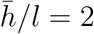, where *l* is the branch length. The node size entropy is

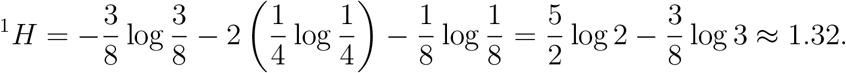

The tree of Figure 7d has the same node sizes and therefore the same ^1^*H* value as the previous tree. It also has the same relative effective height, as

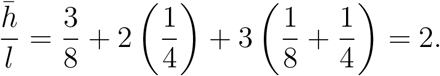

For the tree of Figure 7e, since proportional node sizes must sum to unity we have

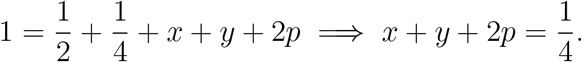

For this tree to have the same 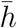 and ^1^*H* values as the four-leaf trees we additionally require

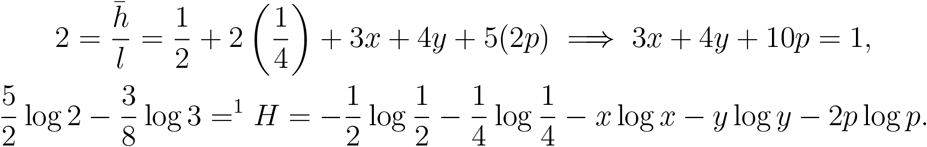

The first two equations together imply *x* = 2*p* and 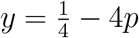 and ^1^*H*. After substituting these results into the third equation we obtain the numerical solution *p* ≈ 0.026606. Since all three trees have identical values of *m*, 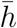 and ^1^*H*, the leafy tree identify implies that they have equal ^1^*D*_*N*_ values and equal balance:

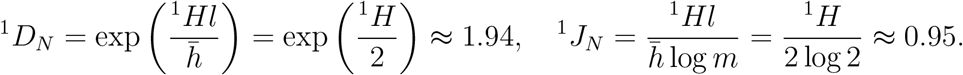

